# SIV-specific neutralizing antibody induction following selection of a PI3K drive-attenuated *nef* variant

**DOI:** 10.1101/2023.04.19.537602

**Authors:** Hiroyuki Yamamoto, Tetsuro Matano

## Abstract

HIV and simian immunodeficiency virus (SIV) infections are known for impaired neutralizing antibody (NAb) responses. While sequential virus-host B cell interaction appears to be basally required for NAb induction, driver molecular signatures predisposing to NAb induction still remain largely unknown. Here we describe SIV-specific NAb induction following a virus-host interplay decreasing aberrant viral drive of phosphoinositide 3-kinase (PI3K). Screening of seventy difficult-to-neutralize SIV_mac239_-infected macaques found nine NAb-inducing animals, with seven selecting for a specific CD8^+^ T-cell escape mutation in viral *nef* before NAb induction. This Nef-G63E mutation reduced excess Nef interaction-mediated drive of B-cell maturation-limiting PI3K/mammalian target of rapamycin complex 2 (mTORC2). *In vivo* imaging cytometry depicted preferential Nef perturbation of cognate Envelope-specific B cells, suggestive of polarized contact-dependent Nef transfer and corroborating cognate B-cell maturation post-mutant selection up to NAb induction. Results collectively exemplify a NAb induction pattern extrinsically reciprocal to human PI3K gain-of-function antibody-dysregulating disease, and indicate that harnessing the PI3K/mTORC2 axis may facilitate NAb induction against difficult-to-neutralize viruses including HIV/SIV.

## Introduction

Virus-specific neutralizing antibody (NAb) responses by B cells are induced by an intricate cooperation of adaptive immune cells (Kumar et al., 2010; Shulman et al., 2014; Giltin et al., 2014; Wang et al., 2014) and often play a central role in clearance of acute viral infections (Junt et al., 2007). In contrast, persistence-prone viruses such as human immunodeficiency virus type 1 (HIV-1), simian immunodeficiency virus (SIV) and lymphocytic choriomeningitis virus (LCMV) variously equip themselves with B cell/antibody-inhibitory countermeasures (Moir et al., 2001; Mattapallil et al., 2005; Sommerstein et al., 2015; Sammicheli et al., 2016; Fallet et al., 2016; Mason et al., 2016), impairing NAb induction (Levesque et al.,2009; Mikell et al., 2011). These viruses successfully suppress elicitation of potent NAb responses, especially in acute infection (Hunziker et al., 2003; Tomaras et al., 2008), and establish viral persistence, posing considerable challenge for developing protective strategies. In particular, HIV and SIV establish early a large body of infection *in vivo* from early on with a distinct host genome-integrating retroviral life cycle (Whitney et al., 2014). In addition, these lentiviruses are unique in launching a matrix of host immune-perturbing interactions mainly by their six remarkably pleiotropic accessory viral proteins, which optimally fuels viral pathogenesis (Kirchhoff et al., 1995; Sheehy et al., 2002; Harris et al., 2003; Schindler et al., 2006; Neil et al., 2008; Zhang et al., 2009; Laguette et al., 2011; Yamada et al., 2018; Joas et al., 2018; Langer et al., 2019; Yan et al., 2019; Joas et al., 2020; Khan et al., 2020; Volcic et al., 2020; Reuschl et al., 2022). A body of evidence has depicted this in the last decades, its entity, including focal mechanisms of humoral immune perturbation in HIV/SIV infection, remains elusive to date.

Adverse virus-host interactions in HIV/SIV infection leads to a detrimental consequence of the absence of acute-phase endogenous NAb responses. Contrasting this, we and others have previously described in *in vivo* experimental models that passive NAb infusion in the acute phase can trigger an endogenous T-cell synergism resulting in robust control of SIV/SHIV (simian/human immunodeficiency virus) (Haigwood et al., 1996; Yamamoto et al., 2007; Ng et al., 2010; Iseda et al., 2016; Nishimura et al., 2017). This indicates that virus-specific NAbs not only confer sterile protection but also can evoke T-cell-mediated non-sterile viral control, suggesting the importance of endogenous NAb responses supported by humoral-cellular response synergisms, during an optimal time frame. Therefore, identifying mechanisms driving NAb induction against such viruses is an important step to eventually design NAb-based HIV control strategies.

One approach that can provide important insights into this goal is analysis of *in vivo* immunological events linked with NAb induction against difficult-to-neutralize SIVs in a non-human primate model. Various *in vivo* signatures of HIV-specific NAb induction, such as antibody-NAb coevolution (Moore et al., 2012), autoimmune-driven induction (Moody et al., 2016) and natural killer cell-related host polymorphisms (Bradley et al., 2018) have been reported to date. The broad range of contributing factors collectively, and interestingly, indicates that pathways to NAb induction against difficult-to-neutralize viruses including HIV/SIV are redundant, and may potentially involve as-yet-unknown mechanisms driving NAb induction. For example, the neutralization resistance of certain SIV strains does not appear to be explained by any of the aforementioned, posing SIV models as attractive tools to analyze NAb induction mechanisms.

In the present study, we examined virus-specific antibody responses in rhesus macaques infected with a highly difficult-to-neutralize SIV strain, SIV_mac239_. This virus is pathogenic in rhesus macaques causing simian AIDS across a broad range of geographical origin of macaques (Cumont et al., 2008). Macaques infected with SIV_mac239_ show persistent viremia and generally lack NAb responses throughout infection (Kestler et al., 1991; Nomura et al., 2012). In this study, a large-scale screening of SIV_mac239_-infected Burmese rhesus macaques for up to 100 weeks identified a subgroup inducing NAbs in the chronic phase. Interestingly, before NAb induction, these animals commonly selected for a specific CD8^+^ cytotoxic T lymphocyte (CTL) escape mutation in the viral Nef-coding gene. Compared with wild-type (WT) Nef, this mutant Nef manifested a decrease in aberrant interaction with phosphoinositide 3-kinase (PI3K)/mammalian target of rapamycin complex 2 (mTORC2), resulting in decreased downstream hyperactivation of the canonical B-cell negative regulator Akt (Omori et al., 2006; Limon et al., 2014). Machine learning-assisted imaging cytometry revealed that Nef preferentially targets Env-specific B cells *in vivo*. Furthermore, the NAb induction was linked with sustained Env-specific B-cell responses after or during the mutant Nef selection. Thus, NAb induction in SIV_mac239_-infected hosts conceivably involves a functional boosting of B cells that is phenotypically reciprocal to a recently found human PI3K gain-of-function and antibody-dysregulating inborn error of immunity (IEI), activated PI3 kinase delta syndrome (APDS) (Angulo et al., 2013; Lucas et al., 2014). Our results suggest that intervening PI3K/mTORC2 signaling can potentially result in harnessing NAb induction against difficult-to-neutralize viruses.

## Results

### Identification of macaques inducing SIV_mac239_-neutralizing antibodies

We performed a retrospective antibody profile screening in rhesus macaques infected with NAb-resistant SIV_mac239_ (*n* = 70) (**Figure 1A**) and identified a group of animals inducing anti-SIV_mac239_ NAb responses (*n* = 9), which were subjected to characterization. These NAb inducers showed persistent viremia with no significant difference in viral loads compared with a subgroup of NAb non-inducers (*n* = 19) that were previously profile-clarified, naïve and major histocompatibility complex class I (MHC-I) haplotype-balanced (used for comparison hereafter) (**Figure 1B** and **Figure 1-figure supplement 1**). Plasma SIV_mac239_-NAb titers measured by 10 TCID_50_ SIV_mac239_ virus-killing assay showed an average maximum titer of 1:16, being induced at an average of 48 weeks post-infection (p.i.) (**Figure 1C**). Two of the nine NAb inducers showed detectable NAb responses by 24 weeks p.i., while the remaining seven induced NAbs after 30 weeks p.i. Anti-SIV_mac239_ neutralizing activity was confirmed in immunoglobulin G (IgG) purified from plasma of these NAb inducers (**Figure 1-figure supplement 2**). SIV Env-binding IgGs developed from early infection in both NAb inducers and non-rapid-progressing NAb non-inducers, the latter differing from rapid progressors known to manifest serological failure (Hirsch et al., 2004; Nakane et al., 2013) (**Figure 1-figure supplement 3A**). Titers of Env-binding IgG were not higher but rather lower at year 1 p.i. in the NAb inducers (**Figure 1-figure supplement 3B** and **3C**). This was consistent with other reports (Havenar-Daughton et al., 2016) and differing with gross B-cell enhancement in Nef-deleted SIV infection with enhanced anti-SIV binding antibody titers accompanying marginal neutralization (Adnan et al., 2016). NAb inducers and non-inducers showed similar patterns of variations in viral *env* sequences (Burns et al., 1993), mainly in variable regions 1, 2 and 4 (V1, V2 and V4) (**Figure 1-figure supplement 4**).

**Figure 1.**
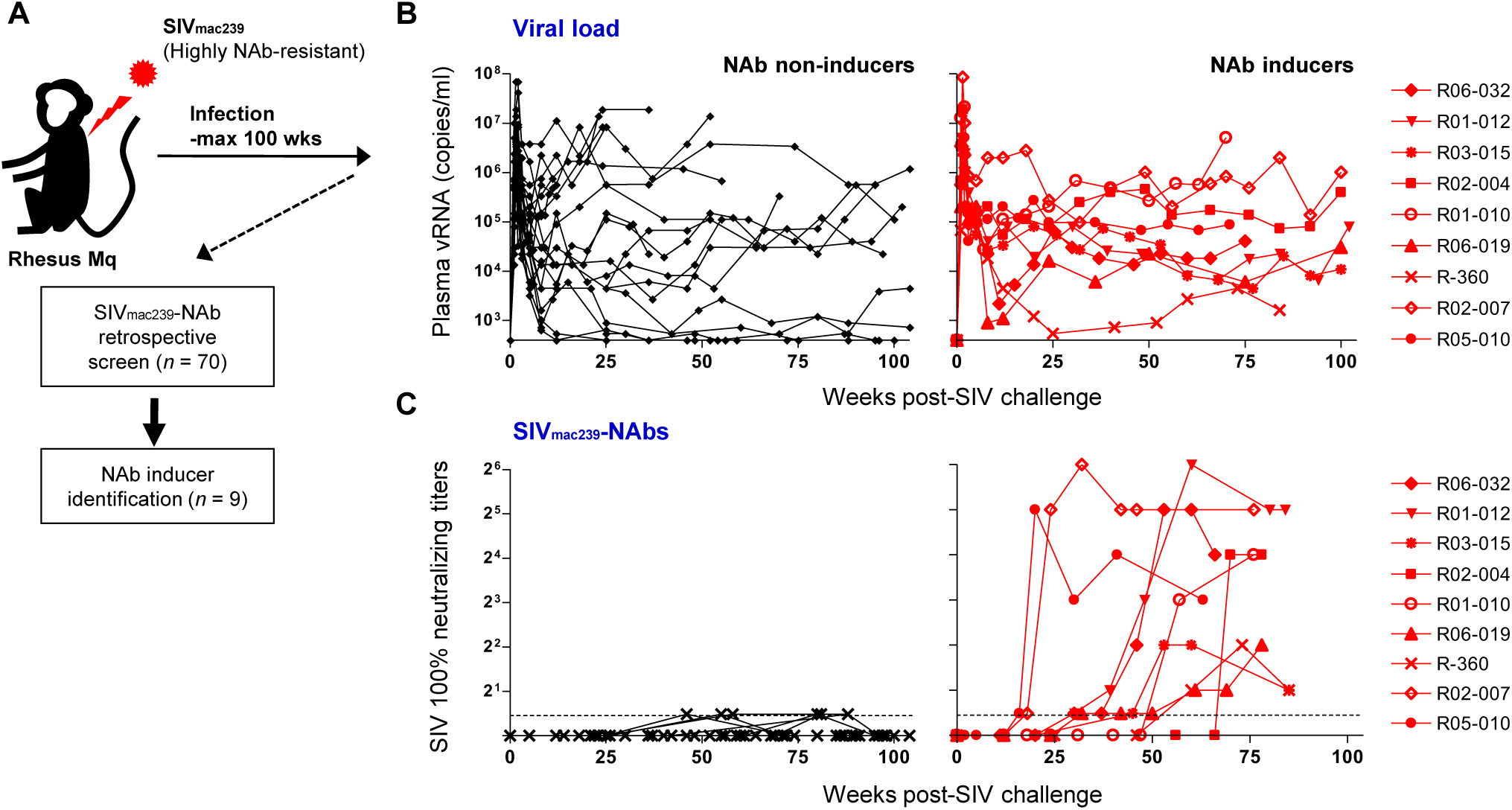
NAb induction against NAb-resistant SIV_mac239._ **(A)** Study design. **(B)** Plasma viral loads (SIV *gag* RNA copies/ml plasma) in NAb non-inducers (left) and inducers (right). **(C)** Plasma SIV_mac239_ 100% neutralizing end point titers by 10 TCID_50_ killing assay on MT4-R5 cells. Points on the dotted line show marginally NAb-positive results (< 1:2). In some animals, titers were comparable with our reported results using MT4 cells.

### Selection of a viral *nef* mutation (Nef-G63E) precedes chronic-phase SIV_mac239_-specific NAb induction

To explore viral mutations linked to NAb induction, we assessed viral nonsynonymous polymorphisms outside *env*. Strikingly, we found selection of a viral genome mutation resulting in G (glycine)-to-E (glutamic acid) substitution at residue 63 of Nef (Nef-G63E) in seven of the nine NAb inducers (**Figure 2A**). Two inducers that did not select Nef-G63E were the early inducers detected for NAb positivity before 24 weeks p.i. This 63rd residue lies in the unstructured N-terminus of Nef, flanked by two α-helices conserved in SIVmac/SIVsmm (**Figure 2B**). This region is generally polymorphic among HIV-1/HIV-2/SIV, occasionally being deleted in laboratory and primary isolate HIV-1 strains (Geyer et al., 2001). AlphaFold2-based structure prediction did not derive palpable change except for low-probability disruption of the alpha helices (data not shown). This mutation was found only in two of the nineteen NAb non-inducers, including one rapid progressor (**Figure 1-figure supplement 1**) and selection was significantly enriched in the NAb inducers as compared to the nineteen non-inducers (**Figure 2C**). Analysis of Nef-G63E mutation frequencies in plasma viruses showed that Nef-G63E selection preceded or at least paralleled NAb induction (**Figure 2D**).

**Figure 2.**
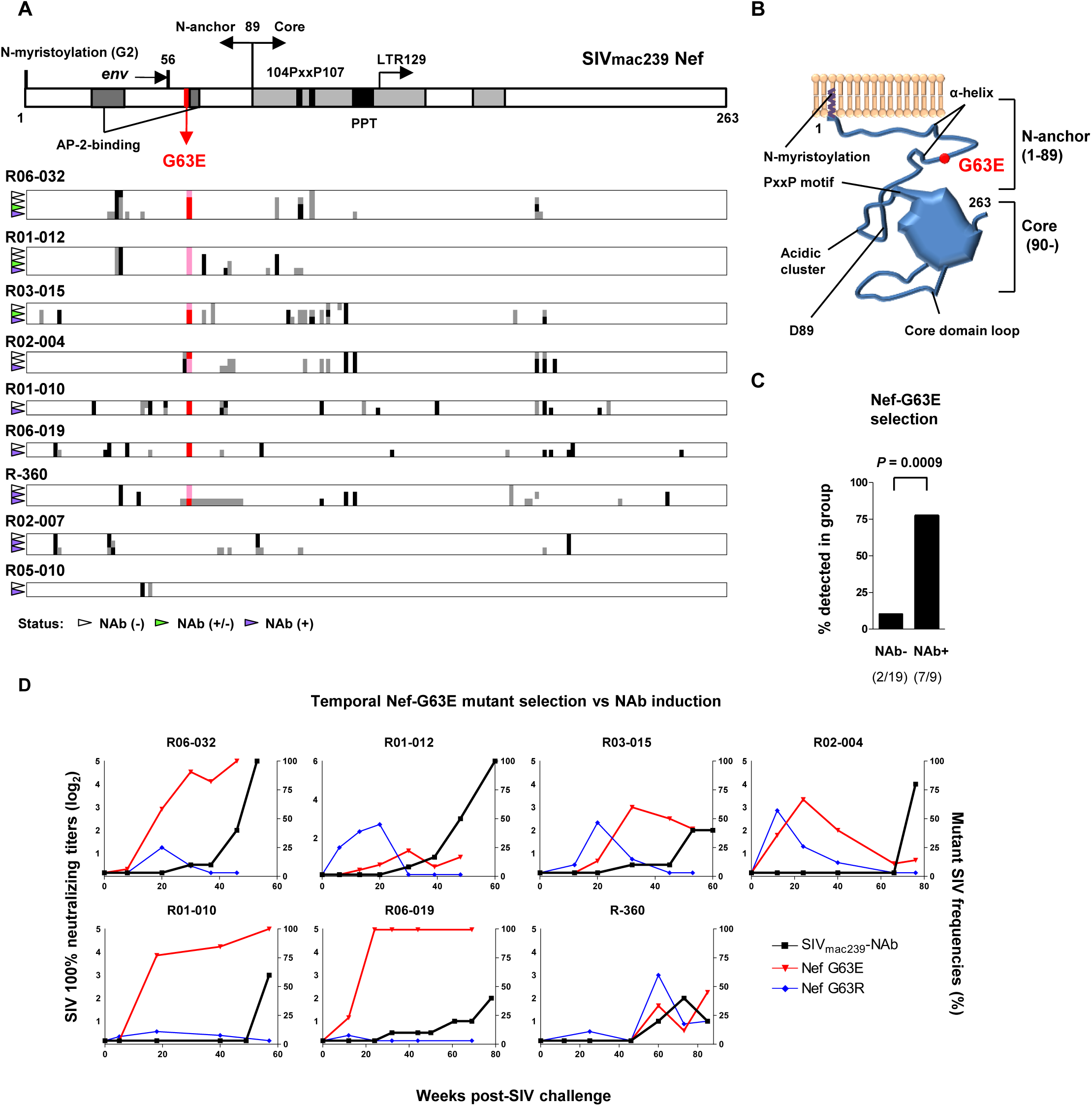
Selection of a viral *nef* mutation (Nef-G63E) before NAb induction. **(A)** Viral *nef* mutations in NAb inducers. A linear schema indicating Nef functional domains is aligned above. Mutations at timepoints with NAb^-^ (indicated by white wedge; mostly before 6 months p.i.), NAb^+/-^ (green wedge) and NAb^+^) (purple wedge; mostly after 1 year) are shown in individual animals. Black and dark gray represent dominant and subdominant mutations (or deletion in R-360) by direct sequencing, respectively. Red and pink indicate dominant and subdominant G63E detection, respectively. **(B)** Schema of SIV_mac239_ Nef structure and Nef-G63E mutation orientation. **(C)** Comparison of frequencies of macaques having Nef-G63E in plasma viruses between NAb non-inducers and inducers. Compared by Fisher’s exact test. **(D)** Temporal relationship of Nef-G63E frequencies in plasma virus and NAb induction. Black boxes (left Y axis) show log_2_ NAb titers; red triangles and blue diamonds (right Y axis) show percentage of G63E and G63R mutations detected by subcloning (15 clones/point on average), respectively.

### Nef-G63E is a CD8^+^ T-cell escape mutation

To explore the mechanistic link between Nef-G63E selection and NAb induction, first we assessed whether CD8^+^ T-cell responses target this region. We found that these seven NAb inducers selecting this mutation elicited CD8^+^ T-cell responses specific for a 9-mer peptide Nef_62–70_ QW9 (QGQYMNTPW) (**Figure 3A**). This Nef_62–70_ QW9 epitope is restricted by MHC-I molecules Mamu-B*004:01 and Mamu-B*039:01 (Evans et al., 1999; Sette et al., 2012). Possession of these accounted for six cases of Nef-G63E selection (**Figure 3A**), and the remaining one animal possessed Mamu-A1*032:02 predicted to bind to Nef_63–70_ GW8 (NetMHCpan). Ten of nineteen NAb non-inducers also had either of these alleles (**Figure 1-figure supplement 1**). This did not significantly differ with the NAb inducer group (*P* = 0.25 by Fisher’s exact test, data not shown), indicating that NAb induction was not simply linked with possession of these MHC-I genotypes but instead required furthermore specific selection of the Nef-G63E mutation (**Figure 2C**). Nef-G63E was confirmed to be an escape mutation from Nef_62-70_-specific CD8^+^ T-cell responses (**Figure 3B**). NAb non-inducers possessing these alleles elicited little or no Nef_62-70_-specific CD8^+^ T-cell responses (**Figure 3-figure supplement 1A**), suggesting that *in vivo* selection and fixation of this Nef-G63E SIV was indeed Nef_62-70_-specific CD8^+^ T cell-dependent. Replication of SIV carrying the Nef-G63E mutation was comparable with WT on a cynomolgus macaque HSC-F CD4^+^ T-cell line (Akari et al., 1996), excluding the possibility that the mutation critically impairs viral replication (**Figure 3-figure supplement 1B**), and plasma viral loads were comparable between Nef-G63E mutant-selecting NAb inducers versus non-inducers (**Figure 3-figure supplement 1C**). These results indicate NAb induction following *in vivo* selection and fixation of the CD8^+^ T-cell escape *nef* mutation, Nef-G63E under viral persistence.

**Figure 3.**
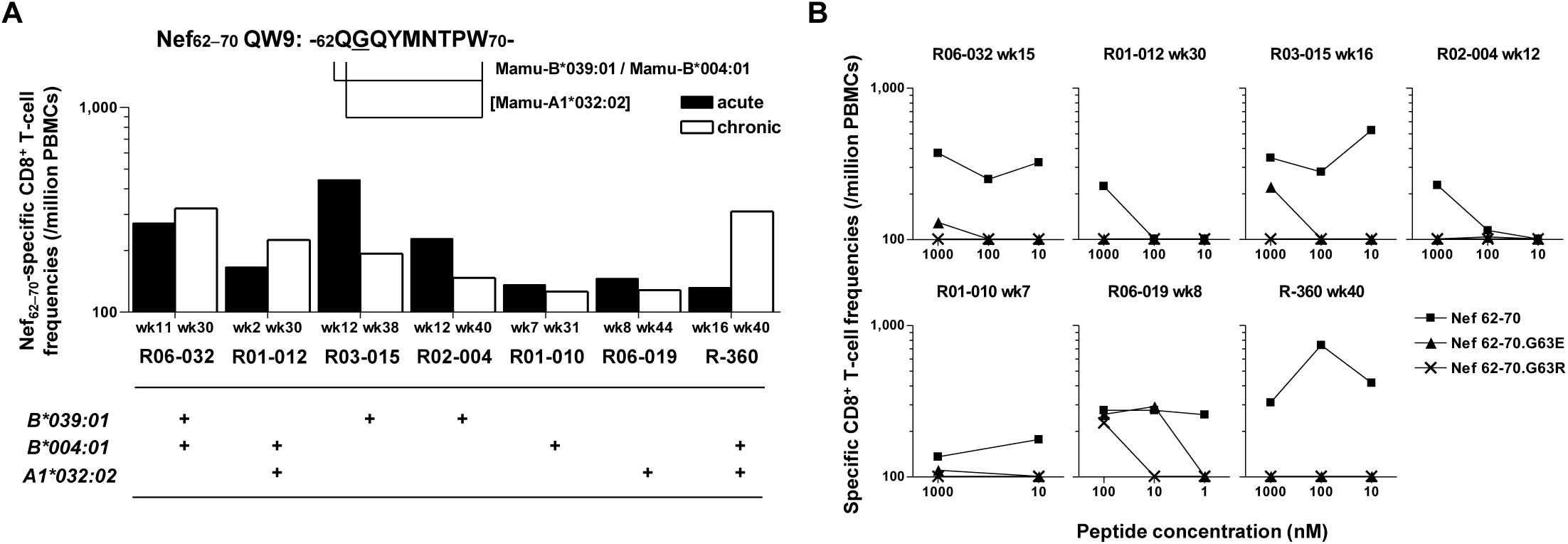
Nef-G63E is a CD8^+^ T-cell escape mutation. **(A)** Nef_62–70_ QW9-specific CD8^+^ T-cell frequencies and related MHC-I alleles in seven NAb inducers selecting Nef-G63E. Mamu-B*039:01 and Mamu-B*004:01 are known to restrict Nef_62–70_ QW9 epitope and binding of Nef_63–70_ peptide to Mamu-A1*032:02 was predicted. **(B)** CD8^+^ T-cell responses specific to wild-type (WT) Nef_62–70_ or mutant Nef_62–70_.G63E or Nef_62–70_.G63R peptides.

### G63E mutation reduces aberrant Nef interaction-mediated drive of PI3K/mTORC2

We next focused on the functional phenotype of Nef-G63E mutant SIV in infected cells. An essence of host perturbation by Nef is its wide-spectrum molecular downregulation, ultimately facilitating viral replication (Schindler et al., 2004; Zhang et al., 2009). To evaluate possible amelioration of this property, we compared downregulation of major targets CD3, CD4, MHC-I, CXCR4 and BST-2 (Jia et al., 2009) in infected HSC-F cells. Nef-G63E mutation did not confer notable change compared with WT [*P* = not significant (*n.s*.) for all molecules, data not shown], implicating other non-canonical changes (**Figure 4A**). One report suggested Nef to drive macrophage production of soluble ferritin and perturb B cells, with its plasma level correlating with viremia (Swingler et al., 2008). Here, plasma ferritin levels showed no differences (**Figure 4-figure supplement 1A**) and viral loads were comparable as aforementioned between Nef-G63E-selecting NAb inducers and non-inducers, arguing against gross involvement of ferritin as well as other viral replication-related Nef phenotypes at least in this model.

**Figure 4.**
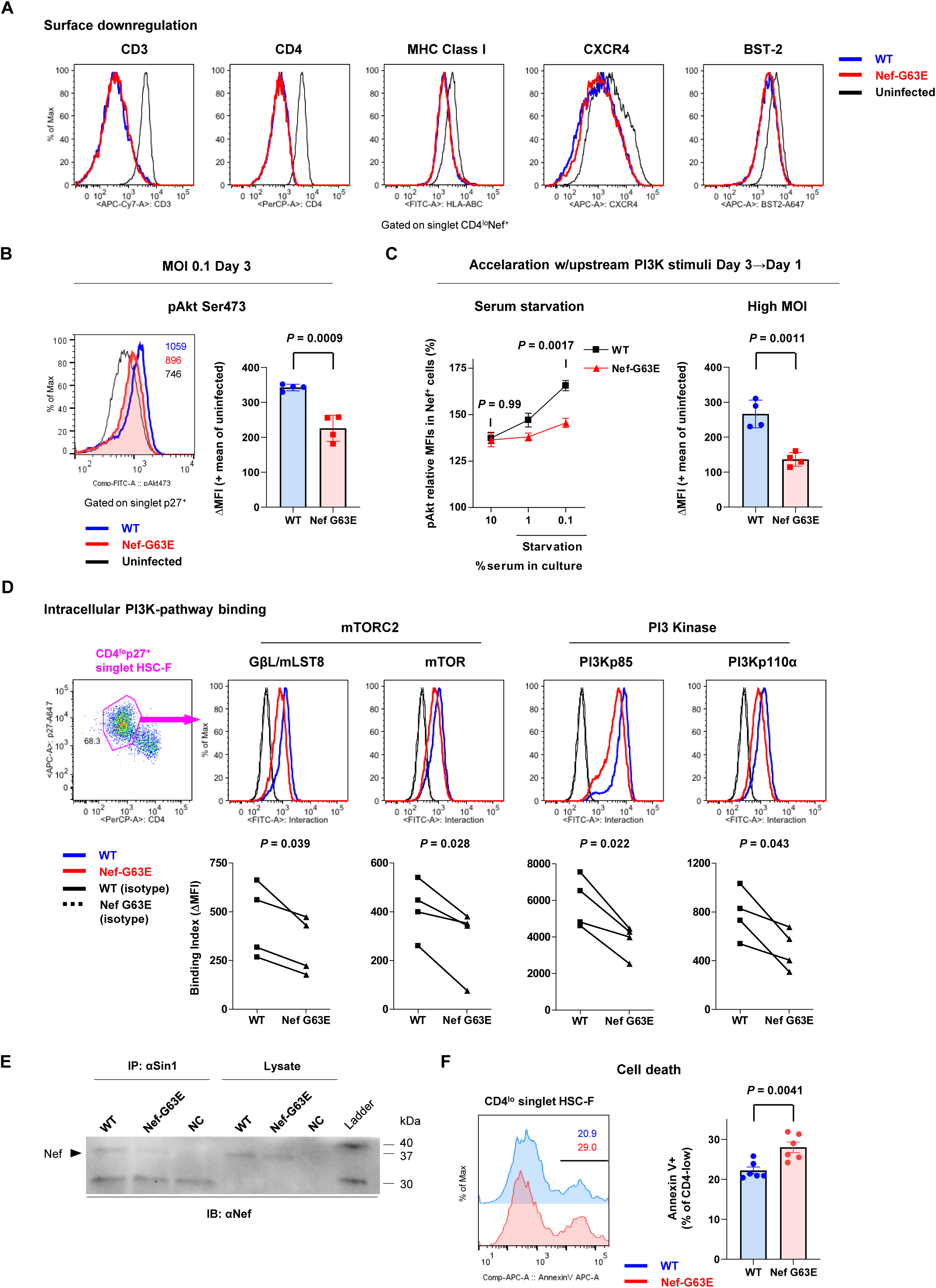
Nef-G63E mutation reduces PI3K/mTORC2 binding and pAkt drive. **(A)** Representative surface expression level histograms of CD3, CD4, MHC class I, CXCR4 and BST-2 in CD4^lo^Nef^+^ subpopulations after WT or Nef-G63E mutant SIV infection at multiplicity of infection (MOI) 0.1 on HSC-F cells. **(B)** Left: representative histograms of relative pAkt serine (Ser) 473 levels in p27^+^ subpopulations after WT or Nef-G63E mutant SIV infection at MOI 0.1 on HSC-F cells. Numbers show pAkt Ser473 mean fluorescence intensities (MFIs) for each. Right: Deviation of pAkt Ser473 MFIs in p27^+^ HSC-F cells compared with mean MFI of uninfected cells. Compared by unpaired t test. **(C)** Left: Relative pAkt Ser473 levels (normalized to mean MFI of uninfected controls) in Nef^+^ HSC-F cells assessed for serum starvation (MOI 0.2, 1 day p.i.). Adjusted *P* values show results of comparison via Sidak’s post-hoc test of 2-way ANOVA (C, left). Right: Deviation of pAkt Ser473 MFIs in Nef^+^ HSC-F cells assessed for high-MOI infection (MOI 5, 1-day p.i.). Compared by unpaired t test. **(D)** Proximity ligation assay (PLA) of Nef binding with mTOR, GβL/mLST8, PI3Kp85 and PI3Kp110α. MFI-based binding index was calculated as (anti-Nef/anti-partner) - (isotype/anti- partner) - (anti-Nef/isotype) + (isotype/isotype). Histograms for samples and isotype/isotype are representatively shown. Differences in MFI binding indexes can be enhanced compared with comparison of raw MFIs. Compared by paired t tests. **(E)** Sin1 co-immunoprecipitation analysis of WT versus Nef-G63E mutant SIV-infected HSC-F cells (infected at MOI 0.05, day 3). Immunoblotting of Nef (37 kDa) in whole cell lysates (lanes 4-6) and anti-Sin1 antibody immunoprecipitates (lanes 1-3) are shown. **(F)** Cell death frequencies of SIV-infected cells measured by Annexin V positivity (% of CD4^lo^) (infected at MOI 0.1, day 3). Compared by unpaired t test. Data represent one of two [(A); (B); (C, right)] independent experiments in quadruplicate, four independent experiments performed in triplicate [(C, left)], four independent single-well comparison experiments pooled for statistical analysis (D), two experiments (E) or two experiments performed with six wells/control (F). Bars: mean ± SD (B, C right), mean ± SEM (C left, F).

Next, we elucidated immunosignaling-modulating properties of Nef-G63E SIV. Akt is known as the predominant immune-intrinsic negative brake of B-cell maturation and AFC/antibody responses *in vivo* (Omori et al., 2006; Limon et al., 2014; Fruman et al., 1999; Ray et al., 2015; Sander et al., 2015). Thus, we analyzed Akt phosphorylation on day 3 after Nef-G63E SIV infection at low multiplicity of infection (MOI). This analysis revealed that its serine 473-phosphorylated form (pAkt Ser473), non-canonically known for Nef-mediated upregulation (Kumar et al., 2016), was significantly lower in Nef-G63E mutant-infected cells compared with WT (**Figure 4B**). The difference observed here was more pronounced than histogram deviation levels in PI3K gain-of-function mice with full recapitulation of B-cell/antibody-dysregulating human APDS disease phenotype (Avery et al., 2018). In contrast, the threonine 308-phosphorylated Akt (pAkt Thr308) level remained unaffected (**Figure 4-figure supplement 1B**).

Interestingly, external PI3K stimuli by serum starvation (Kennedy et al., 1997) accelerated phenotype appearance from day 3 to day 1 post-infection (**Figure 4C**, left). A similar trend was obtained by short-term PI3K stimulation with interferon-γ (Nguyen et al., 2001), interleukin-2 (Marzec et al., 2008) and SIV Env (François & Klotman, 2003) (**Figure 4-figure supplement 1C**). A high-MOI SIV infection, comprising higher initial concentration of extracellular Env stimuli, also accelerated phenotype appearance from day 3 to day 1 post-infection, with stronger pAkt reduction (**Figure 4C**, right). These data indicate that the Akt-inhibitory G63E mutant Nef phenotype is PI3K stimuli-dependent. Transcriptome analysis signatured decrease in PI3K-pAkt-FoxO1-related genes in mutant SIV infection (**Figure 4-figure supplement 1D**). Given that Akt is also a survival mediator, Nef-G63E mutation may be Th-cytopathic and decrease Th dysfunctional expansion (Baumjohann et al., 2013), as had been observed in LCMV strain WE (Recher et al., 2004) and chronic HIV/SIV (Gray et al., 2011; Lindqvist et al., 2012; Petrovas et al., 2012) infections. Here, reduced chronic-phase peripheral CXCR3^-^CXCR5^+^PD-1^+^ memory follicular Th (Tfh), linked with antibody cross-reactivity in one cohort (Locci et al., 2013), was somewhat in line with this notion (**Figure 4-figure supplement 1E** and **1F**). We additionally examined another CTL escape mutant Nef-G63R, selected with marginal statistical significance primarily in early stage (**Figure 4-figure supplement 2A**). This mutant did not associate with NAb induction in one control animal (R06-034) even upon preferential selection (**Figure 4-figure supplement 2B**) and Nef-G63R mutant was not similarly decreased in pAkt drive (**Figure 4-figure supplement 2C**), implicating that the Nef-G63E mutation was more tightly linked with NAb induction in terms of an immunosignaling phenotype.

We further explored molecular traits of this decreased Akt hyperactivation. We reasoned that comparing endogenous Nef binding patterns would be adequate and analyzed SIV-infected HSC-F cells with a flow cytometry-based proximity ligation assay (PLA) (Leuchowius et al., 2009; Avin et al., 2017). We found that this Nef-G63E mutation causes significant decrease in Nef binding to PI3K p85/p110α and downstream mTORC2 components mTOR (Sarbassov et al., 2005) and GβL/mLST8 (Kim et al., 2003) in the CD4^lo^-SIV Gag p27^+^ infected population (**Figure 4D**). Results were biochemically confirmed by a decrease in G63E-mutated Nef binding to the mTORC2-intrinsic cofactor Sin1 in coimmunoprecipitation analysis of infected HSC-F cells (**Figure 4E**). Collectively, the Nef-G63E mutation attenuates PI3K/mTORC2 signaling driven by aberrant Nef bridging, explaining decreased Akt Ser473 phosphorylation by mTORC2. These properties were in keeping with more pronounced apoptosis of CD4-downregulated/infected HSC-F cells upon Nef-G63E SIV infection (**Figure 4F**). Thus, Nef-G63E SIV is a mutant virus decreased in aberrant interaction/drive of B-cell-inhibitory PI3K/mTORC2 signaling, manifesting a molecular signature reciprocal to human APDS (Angulo et al., 2013; Lucas et al., 2014; Avery et al., 2018).

### Targeting of lymph node Env-specific B cells by Nef *in vivo*

Next, we analyzed *in vivo* targeting of virus-specific B cells by Nef in lymph nodes to explore the potential B-cell-intrinsic influence of the Nef-G63E phenotype. Previously suggested influence of soluble Nef itself (Qiao et al., 2006) and/or related host factors (Swingler et al., 2008) may derive generalized negative influence on B cells. However, SIV antigen-specific binding antibody responses were rather decreased in NAb inducers (**Figure 1-figure supplement 1**), with comparable viral loads (**Figure 3-figure supplement 1C**) and ferritin levels (**Figure 4-figure supplement 1A**), differing from the aforementioned literature stating positive correlation between the three. We surmised that some targeted Nef intrusion against Env-specific B cells may be occurring and that the decreased aberrant drive of Nef-PI3K/mTORC2 may result in their enhanced maturation in lymph nodes.

While reports (Xu et al., 2009) histologically proposed Nef B-cell transfer, quantitative traits, e.g., invasion frequency and influence on virus-specific B cells, have remained unvisited. One of the reasons is that intracellular Nef staining is dim (particularly for B cell-acquired Nef) and difficult to examine by conventional flow cytometry. To circumvent this issue, defining staining cutline using single-cell images potentially overcomes confounding technical hurdles, such as high Nef false-staining signals owing to pre-permeation rupture and post-permeation processing that derives biologically discontinuous staining and/or batch-inflated signals. Thus, we reasoned that at the expense of spatial information, sophisticating imaging cytometry would best visualize Nef-mediated B-cell perturbation *in vivo*. We analyzed lymph node B cells with Image Stream X MKII, with high-power (> 10-fold) antigen detection [e.g., molecules of equivalent soluble fluorochrome (MESF) 5 *vs.* MESF 80 in an average flow cytometer for FITC detection] ideal for detection/single-cell verification.

We designed a triple noise cancellation strategy to overcome the issues stated above. Firstly, amine reactive dye staining (Perfetto et al., 2006) gated out B cells being Nef-positive due to pre-experimental membrane damage (**Figure 5-figure supplement 1A**, left). Next, we deployed secondary quantitative parameters of Nef signals deriving from each pixel of the images. Generation of a gray-level cooccurrence matrix (GLCM) (Haralick et al., 1973), which computes adjacent signal deviation as their frequencies on a transpose matrix, enables calculation of a variety of feature values summarizing traits of the whole image. A biological assumption of Nef signal continuity in a true staining suggested that the sum of weighted square variance of GLCM, i.e., the {sum of [square of (value – average signal strength) × frequency of each value occurrence]} would separate natural versus artificial Nef signals. This value, known as Haralick variance, was computed for multiple directions and averaged. The resultant Haralick variance mean (**Figure 5-figure supplement 1A**, X axis) is proportionate with unnaturalness of signals in the Nef channel. Finally, Nef signal intensity threshold (**Figure 5-figure supplement 1A**, Y axis) gated out overtly-stained cells void for true-false verification.

**Figure 5.**
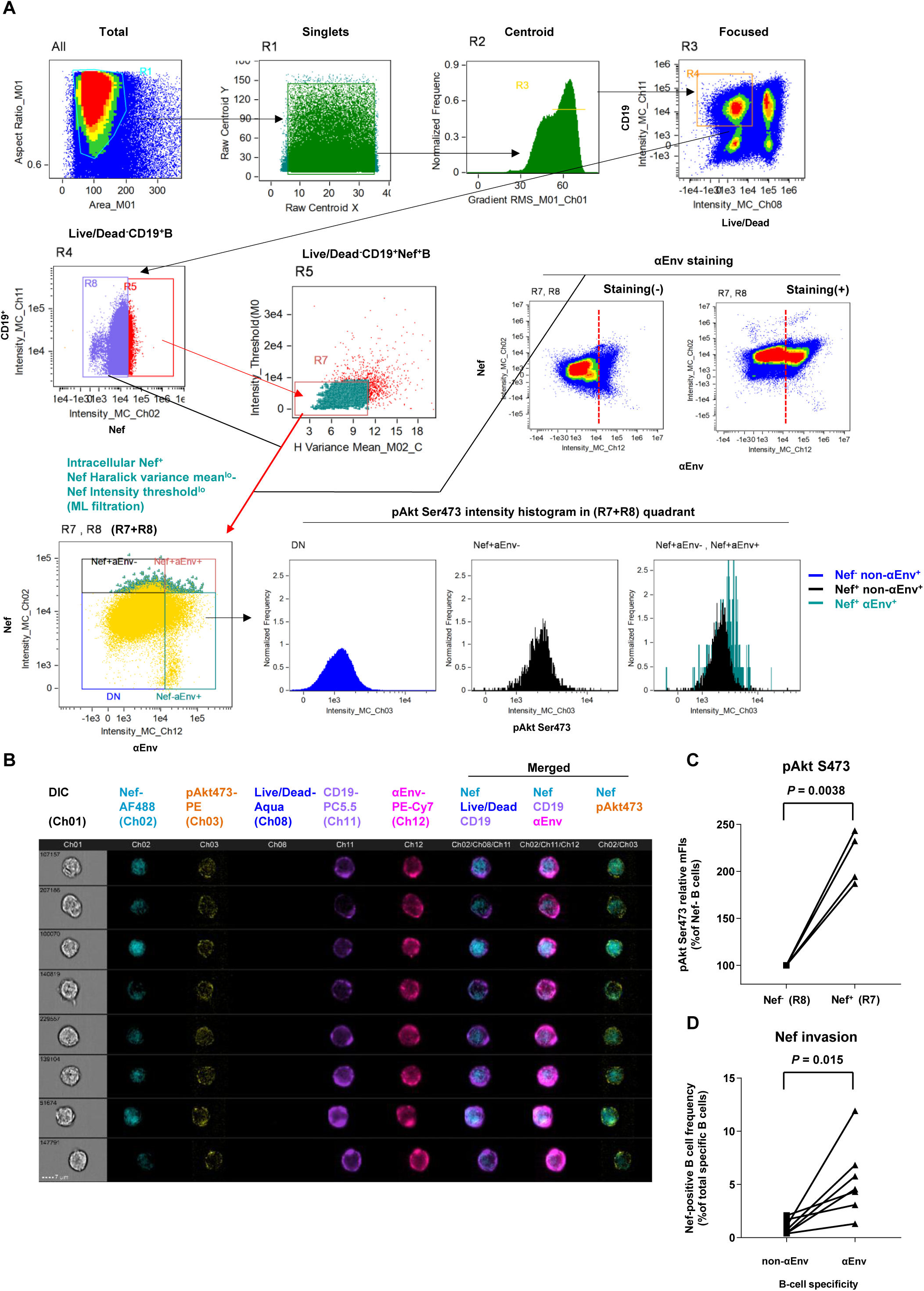
Preferential targeting of lymph node Env-specific B cells by Nef *in vivo*. **(A)** Representative gating of triple noise-cancellation *in vivo* Nef staining in B cells analyzed by ImageStreamX MKII. Pre-experimental damaged cells are first excluded with Live/Dead from focused/centroid/singlet image-acquired CD19^+^ B cells (first lane right/R4 gated on “Focused”). Following Nef^+^ gating (second lane left/R5 gated on R4), a second step of Nef noise cancellation (second lane middle/R7 on R5) comprises double-negative removal of post-experimental stochastic irregular staining building a disparate intracellular staining gradient (X axis, Nef signal pixel Haralick variance mean) and post-experimental batch overt cellular staining (Y axis, Nef signal pixel intensity threshold). This outputs a B-cell population with a fine-textured pericellular Nef^int-lo^ staining, biologically concordant with Nef membrane-anchoring. Probing of anti-Env BCR (αEnv) by recombinant SIV Env (second lane, right) is combined, resulting in a 2-D panel of intracellular Nef versus αEnv for noise-cancelled Nef^+^ B cells (R7) plus Nef^-^ B cells (R8) (third lane left/“R7+R8”). pAkt Ser473 expression (third lane right) and cellular morphology (B) was further analyzed. DN, double-negative. **(B)** Typical images of Nef-transferred Env-specific B cells defined αEnv^+^-intracellular Nef^+^-Nef Haralick variance mean^lo^-Nef Intensity threshold^lo^-Live/Dead^-^-CD19^+^ cells [“Nef^+^aEnv^+^” population of lower left panel in (A), gated on “R7, R8”]. Note the pericellular pAkt Ser473 upregulation in these cells (Ch 03/yellow). Data on inguinal lymph node lymphocytes of macaque R10-007 at week 62 post-SIV_mac239_ infection are shown in (A) and (B). **(C)** Comparison of pAkt Ser473 median fluorescence (medFI) intensity levels in Nef^-^ B cells (R8) versus noise-cancelled Nef^+^ B cells (R7). Analyzed by paired t test. **(D)** Comparison of Nef-positive cell frequencies in non-Env-specific (left) versus Env-specific (right) B cells in lymph nodes of persistently SIV-infected macaques (*n* = 6). Analyzed by paired t test.

These measures excluded B cells with non-specific, strong-signal binary-clustered staining pixels for Nef (deriving a large summated variance: X axis, and/or batch-stained for Nef: Y axis; likely originating from post-experimental membrane damage).

Using this approach, we successfully acquired images of viable Nef^+^CD19^+^ B cells with low Nef signal Haralick variance mean and low Nef signal intensity threshold, with fine-textured gradation of low-to-intermediate Nef staining with continuity from membrane-proximal regions and without sporadic staining speckles (**Figure 5A** and **Figure 5-figure supplement 1**). Scoring segregation of typical void/valid images by a linear discriminant analysis-based machine learning module showed that this gating provides the highest two-dimensional separation (**Figure 5-figure supplement 1**). Combined with visualization of Env-binding B cells, we reproducibly obtained images of Env-specific (Ch 12) CD19^+^ (Ch 11) B cells without membrane ruptures (Ch 08) and showing fine-textured transferred Nef (Ch 02) (**Figure 5B**). Nef invasion upregulated pAkt Ser473 (Ch 03) to a range resembling *in vitro* analysis (**Figure 5C**), demonstrating that Nef-driven aberrant PI3K/mTORC2 signaling does occur in Nef-invaded B cells *in vivo*. Strikingly, lymph node Env-specific B cells showed significantly higher Nef-positive frequencies as compared with batch non-Env-specific B cells (**Figure 5D**). This indicated that infected cell-derived Nef preferentially targets adjacent Env-specific B cells, putatively through its contact/transfer from infected CD4^+^ cells to B cells (Xu et al., 2009; Hashimoto et al., 2016). Thus, the phenotypic change in Nef can directly dictate Env-specific B-cell maturation.

### Cell contact-dependent B-cell invasion from infected cells *in vitro*

To understand cell-intrinsic properties enhancing B-cell Nef acquisition *in vivo*, we addressed in an *in vitro* coculture reconstitution how infected cell-to-B-cell Nef invasion takes place and how it is modulated. We performed imaging cytometry on a twelve-hour coculture of SIV-disseminating HSC-F cells (reaching around 30% Nef-positivity at moment of coculture) with Ramos B cells, with or without modulators added throughout the coculture period (**Figure 6A**, top left). In this coculture, there is no MHC-related interaction known to enhance T-cell/B-cell contact (Wülfing et al., 1998). Machine learning-verified noise cancellation of fragmented signal acquisition (**Figure 6A**, middle left) filtered reproducible images of Nef-positive HSC-F cells directly adhering to Ramos B cells within the doublet population (**Figure 6A**, top to middle right). Importantly, addition of soluble, stimulation-competent antibodies against macaque CD3 enhanced adhesion of Nef^+^ T cells to B cells (**Figure 6A**, bottom center). In these doublets, we readily detected images of polarized Nef protrusion/injection from T cells to B cells (second column, filled wedges) as well as trogocytosed B-cell membranes (third column, open wedge), suggesting dynamic Nef acquisition by B cells. Changing centrifugation strengths upon coculture collection and staining (190 x *g* versus 1,200 x *g* and 600 x *g* versus 1,200 x *g*, respectively) shifted the tendency between enriched Nef^+^ HSC-F-B-cell doublet versus singlet detection via CD3 stimulation, respectively (**Figure 6A**, lower right).

**Figure 6.**
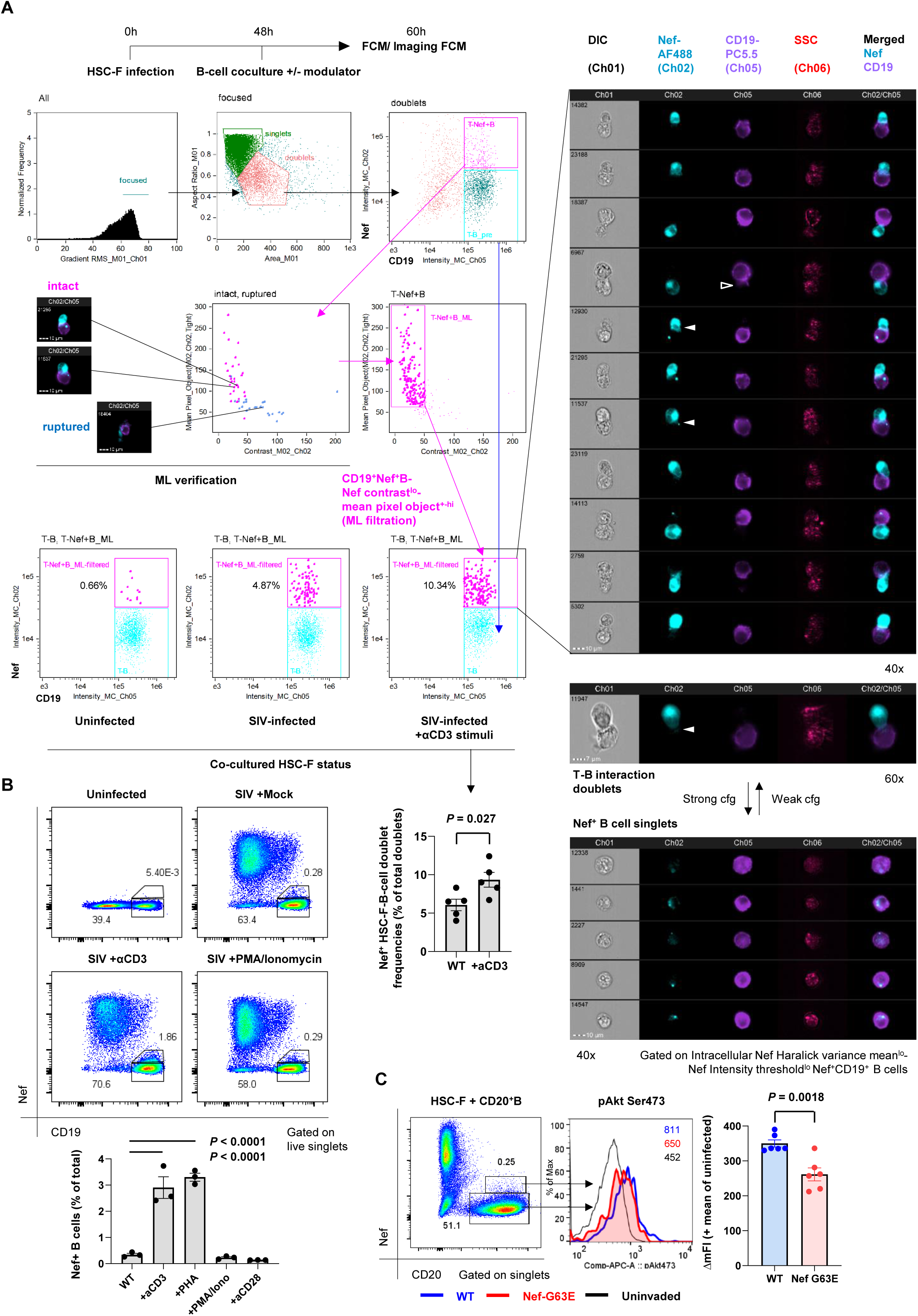
Contact-dependent enhancement of B-cell Nef acquisition in reconstitution. **(A)** Imaging flow cytometric analyses of stimuli-dependent enhancement of infected HSC-F cell adhesion to non-permissive CD19^+^ Ramos B cells with the indicated gating (left). Cocultured doublet cells subjected to machine learning-verified filtration of fragmented signal acquisition (via gating Nef pixel signal contrast^lo^-Nef mean pixel per object^hi^ cells) deriving typical images of infected HSC-F cell (Nef stain shown in pale blue pseudocolor) adhesion to Ramos cells (CD19 stain shown in purple pseudocolor) are shown. Analyzed by unpaired t test for doublet formation in the unstimulated versus anti-CD3 monoclonal antibody (clone FN-18)-stimulated group. **(B)** Flow cytometric analyses of Nef-acquiring live CD19^+^ Ramos B cells upon coculture with SIV-infected HSC-F cells with or without the indicated stimuli. Analyzed by one-way ANOVA with Tukey’s post-hoc multiple comparison tests. **(C)** Representative flow cytometric plot (left), histogram (middle) and pAkt Ser473 signal deviation in WT versus G63E Nef-acquiring CD20^+^ primary B cells (six wells/control). Analyzed by unpaired t test. Data represent pooled data of two (A) or three [(B) and (C)] independent experiments with indicated number of replicates showing similar results.

Substantiated by the acquired images, an enhancement of live singlet Ramos B-cell Nef acquisition (% Nef^+^ in total B cells) was also observed by CD3 stimulation in conventional flow cytometry (**Figure 6B**). There was a similar enhancement when we added phytohemagglutinin (PHA) whereas phorbol-12-myristate-13-acetate (PMA)/ionomycin did not affect Nef acquisition by B cells. In a similar 72-hour static coculture with primary CD20^+^ B cells, we detected recapitulation of the decreased B-cell pAkt Ser473 phosphorylation upon G63E Nef transfer compared to that of WT Nef (**Figure 6C**), indicating that the Nef-G63E mutation can directly alleviate maturation inhibition in preferentially Nef-targeted Env-specific B cells.

### Enhanced Env-specific B-cell responses after PI3K-diminuting mutant selection

Finally, to assess *in vivo* B-cell quality in NAb inducers with Nef-G63E mutant selection, we examined peripheral SIV Env-specific IgG^+^ B-cell responses comprising plasmablasts (PBs) and memory B cells (B_mem_). After excluding lineage-specific cells [T/NK/pro-B cells/monocytes/myeloid dendritic cells (DCs)/plasmacytoid DCs], we defined IgG^+^ PBs as showing: high replication (Ki-67^+^), post-activation (HLA-DR^+^), transcriptional switching for terminal differentiation (IRF4^hi^) and downregulated antigen surface binding (surface [s]Env^lo^-cytoplasmic [Cy]Env^+^) (Nutt et al., 2015; Silveira et al, 2015) (**Figure 7A** and **Figure 7-figure supplement 1**). NAb inducers showed significantly higher Env-specific IgG^+^ PB responses around week 30 p.i., after Nef-G63E selection, compared to those in NAb non-inducers (**Figures 7B** and **7C**). At 1 year, this difference became enhanced as a result of less pronounced decrease in Env-specific PB frequencies in the NAb inducers.

**Figure 7.**
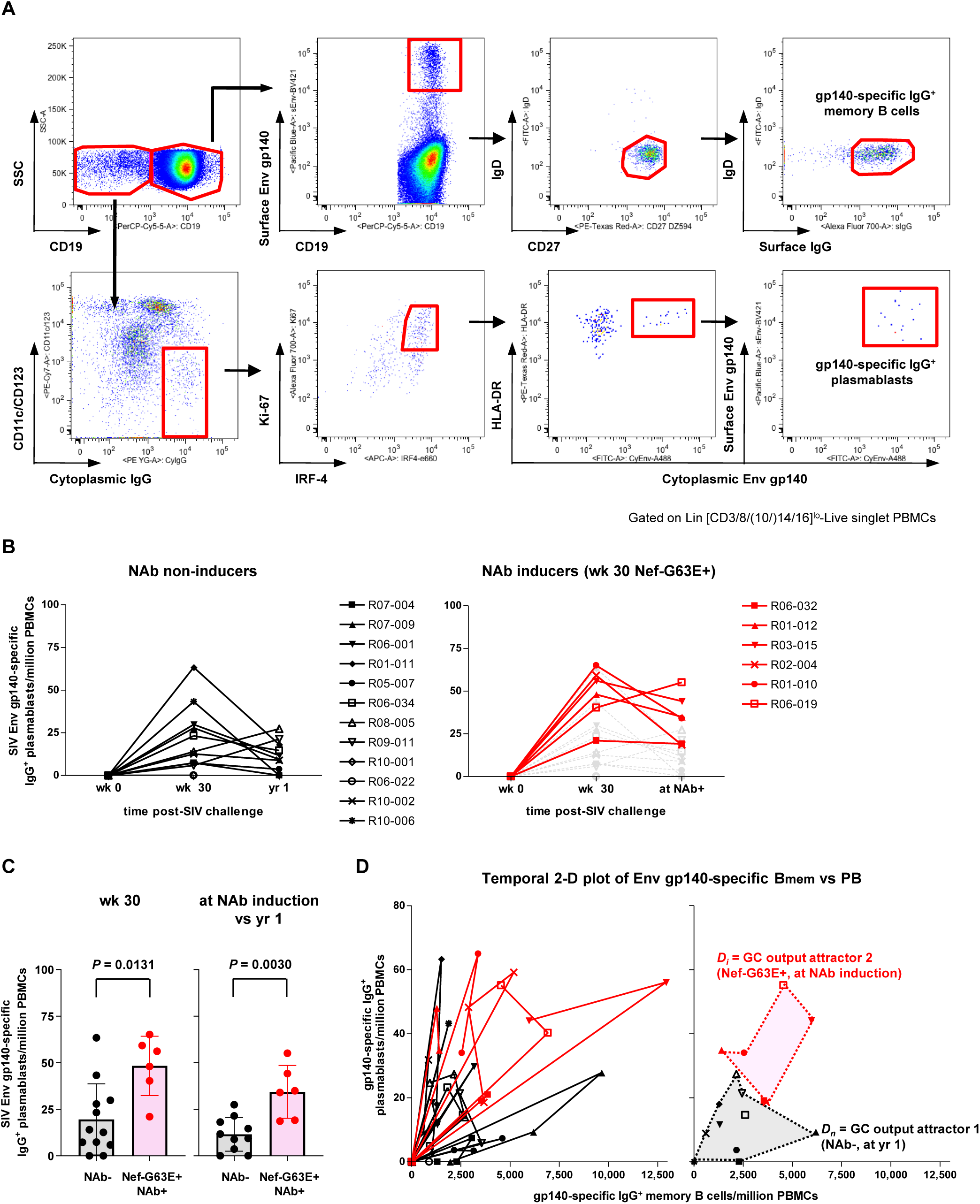
Enhanced SIV Env-specific B-cell output up to NAb induction following Nef-G63E selection. **(A)** Representative gating (R02-004, week 32 p.i.) of SIV Env gp140-specific memory B cells (B_mem_) and plasmablasts (PBs). Two panels for PB gating are shown in lower resolution for visibility. B_mem_ staining was performed separately and gating are merged with PB panels (second row, panels 2–4). For lineage-specific exclusion (first row, panel 4), CD10 was stained independently for B_mem_ (see parenthesis). **(B)** Changes in SIV Env gp140-specific plasmablast (PB) frequencies in viremic NAb non-inducers (left, *n* = 12) and Nef-G63E-selecting NAb inducers (right, *n* = 6). Available samples of twelve viremic NAb non-inducers (including ten with ≥ 1-year survival) and six NAb inducers were tracked. In the right panel, the frequencies in NAb non-inducers are shown in background (gray) for comparison. **(C)** Comparison of Env gp140-specific PB frequencies between NAb non-inducers and inducers by Mann Whitney U tests. In the right, the frequencies at NAb induction were compared with those at year 1 in NAb non-inducers with ≥ 1-year survival (*n* = 10). Bars: mean ± SD. **(D)** Left: vector chart of Env gp140-specific memory B cell (B_mem_) and PB levels. Legends for each animal correspond to the ones in (B). Right: NAb non-inducer vectors empirically define a polygonal GC output attractor area *D_n_* (gray area surrounded with dotted lines) on which they converge by and beyond year 1 p.i. The NAb inducer vectors (shown up to the time of NAb induction) remained outside of *D_n_*. At the moment of NAb induction they converged on a second GC output attractor area *D_i_* (red area surrounded with dotted lines), mutually exclusive with *D_n_* (*P* < 0.0001 by Fisher’s exact test on NAb inducer vector convergence frequency within *D_n_*). Legends for year 1 p.i. in the NAb non-inducers and moment of NAb induction in the NAb inducers are specified.

Simultaneous quantification of Env-specific PB and IgD^-^CD27^+^IgG^+^ B_mem_ responses allowed for an assessment of overall B-cell response quality by pair-wise analysis of Env-specific IgG^+^ B_mem_/PB as a projection of germinal center (GC) output (Zotos et al., 2010) (**Figure 7D**, left). In this two-dimensional vector temporally plotting the frequency of B_mem_ and PBs, vector protrusion towards upper right represents higher gross GC output of antibody-forming cells (AFCs). In NAb non-inducers, all vectors converged on an empirically defined polygonal attractor area *D_n_* (gray area surrounded with dotted lines in **Figure 7D**, right) at year 1 p.i. and beyond, describing that these NAb non-inducers failed to sustain Env-specific GC output. In contrast, the vectors were consistently tracked outside *D_n_* in the NAb inducers with Nef-G63E (**Figure 7D**). At the time of NAb induction, they converged on another upper-right GC output attractor area *D_i_* (red area surrounded with dotted lines in **Figure 7D**, right) that is mutually exclusive with *D_n_* (*P* < 0.0001 by Fisher’s exact test on NAb non-inducer/inducer vector distribution frequency within *D_n_*). These suggest more robust virus-specific IgG^+^ B-cell responses following Nef-G63E CD8^+^ T-cell escape mutant selection, ultimately leading to NAb induction. The enhanced signature of cognate B cells was also an inverted pattern of impaired terminal differentiation of B-cell responses in APDS (Al Qureshah et al., 2021).

Taken together, in the current model, Nef_62-70_-specific CD8^+^ T-cell responses in persistently SIV_mac239_-infected macaques selected for an escape mutant, Nef-G63E, which displays attenuated aberrant Nef binding and ensuing drive of B-cell-inhibitory PI3K/mTORC2. Nef invasion of B cells *in vivo* occurred more preferentially in Env-specific B cells, suggesting diminution in Nef-mediated tonic dysregulation of B cells after mutant selection. These events conceivably predisposed to enhanced Env-specific B-cell responses and subsequent SIV_mac239_-specific NAb induction, altogether, in a manner reciprocal to human APDS-mediated immune dysregulation.

## Discussion

In the present study, we found that in macaques infected with an NAb-resistant SIV, selection of a CD8^+^ T-cell escape *nef* mutant virus, Nef-G63E, precedes NAb induction. As a result, Nef binding-mediated drive of PI3K/mTORC2 in Env-specific B cells becomes attenuated, which in turn unleashes maturation of antiviral B-cell responses *in vivo* to induce NAbs. Importantly, this manifested through a pAkt deviation level more pronounced than WT versus germline *PIK3CD* gain-of-function mutation heterozygote mice recapitulating full human APDS phenotype (Avery et al., 2018). Thus, the current Nef-G63E-associated B-cell/NAb phenotype likely occurs in a “reciprocal APDS-like” manner. This proposition was enhanced based on extension of immune cell-intrinsic Nef influence on cognate B cells (Fig. 5), in addition to infected T cells. This work is, to our knowledge, the first to interlink a PI3K/mTORC2-modulating viral signature and enhanced B-cell/NAb responses in a viral infection model.

A link between viral T-cell escape and consequent immune modulation has been previously explored to some extent. For example, enhanced binding of mutant HIV-1 epitope peptide to inhibitory MHC-I on DCs impairs T cells (Lichterfeld et al., 2007) and decreased CTL-mediated lymph node immunopathology can drive, and not inhibit, the production of LCMV-specific antibodies (Battegay et al., 1993). Our results now evidence a new pattern of NAb responses that are bivalently shaped through viral interactions with both humoral and cellular immunity in AIDS virus infection. SIV_mac239_ infection of macaques possessing MHC-I alleles associated with Nef-G63E mutation can be one unique model to analyze virus-host interaction for B cell maturation leading to NAb induction. The interval between Nef-G63E mutant selection and NAb induction was variable in the NAb inducers, for which we did not obtain a clear explanation. This may be influenced by certain basal competitive balance between humoral versus cellular adaptive immunity, as observed in certain vaccination settings (Querec et al., 2009).

The current Nef-G63E phenotype identified here adds yet another aspect to the wealth of evidence documenting the multifaceted impact of Nef in HIV/SIV infection (Kirchhoff et al., 1995; Gauduin et al., 2006; Stolp et al., 2012). Based on our data, the current phenotype differed from decreased replication-related properties or generalized amelioration in immune impairment such as those of delta-Nef HIV/SIV (Kirchhoff et al., 1995; Johnson et al., 1997; Gauduin et al., 2006; Fukazawa et al., 2012). The SIV Nef N-terminal unstructured region comprising Nef-G63E is not conserved in HIV-1 (Schindler et al., 2004) and beyond this model study it remains to be addressed whether a mutant HIV-1 with a similar immunosignaling-related phenotype can be obtained and how much lentiviral conservation exists for such interactions. Analysis of cognate B-cell maturation on cohort basis (Hau et al., 2022) may potentially assist this approach.

Preferential Nef transfer to Env-specific B cells (**Figure 5**) suggests potential involvement of polarized perturbation in cognate immune cells *in vivo*, which was best depicted for preferential infectivity of HIV-specific CD4^+^ T cells (Douek et al., 2002) and HIV condensation to the interface of dendritic cell-T-cell contacts (McDonald et al., 2003). As cognate MHC class II interactions (Wülfing et al., 1998; Giltin et al., 2014) regularly occur between HIV/SIV-specific CD4^+^ T cells and Env-specific B cells in GCs, dysregulation of Env-specific B cells may well result from hijacking of cognate T-cell/B-cell interactions by the virus. Enhanced T-cell attachment and transfer of Nef to B cells by TCR stimulation (**Figure 6**) may override constraints of infection-mediated CD4^+^ T-cell cytoskeletal impairment (Campbell et al., 2004; Jolly et al., 2004; Stolp et al., 2009) particularly by antagonizing reduced motility (Stolp et al., 2012), resulting in *in vivo* “filtering” of enhanced cognate CD4^+^ T-cell interaction with cognate B cells. Interestingly, Tfh cells that bear trogocytosed CD20 and are positive for viral DNA appear *in vivo* during SIV infection and increase when infection progresses (Samer et al., 2023). This is consistent with our data that suggest occurrence of contact-dependent, polarized interactions facilitating Nef transfer to cognate B cells.

The major *in vivo* readout of this study is autologous neutralization of a highly difficult-to-neutralize SIV strain, which is different from HIV-specific broadly neutralizing antibodies (bNAbs). Importantly, however, obtained signatures of enhanced IgG^+^ Env-specific AFC and B cells (class switch/terminal differentiation) and virus neutralization (hypermutation) in our model are indicative of elevated activity of activation-induced cytidine deaminase (AID), the canonical positive driver of B-cell fate. Thus, we reason that findings in this study have a strong conceptual continuity with bNAb regulation by AID in HIV infection.

Our results extracted an *in vivo* link between a decrease in aberrant Nef-PI3K/mTORC2 interaction and major enhancement in B-cell responses. The relationship is reciprocal to immunogenetic mechanisms of human PI3K gain-of-function mutations resulting in APDS with multiply impaired anti-viral B-cell responses. APDS occurs from germline mutations in leukocyte-intrinsic p110δ catalytic subunit-coding *PIK3CD* or the ubiquitous p85α regulatory subunit-coding *PIK3R1*. In both cases, the mutations cause residue substitutions resulting in class IA PI3K gain-of-function (Dornan et al., 2017). Hyperactivated PI3K interferes with cognate B cells in an immune cell-intrinsic manner, leading to impaired B-cell class switching and resultant susceptibility against various infections (Angulo et al., 2013; Lucas et al., 2014; Coulter et al., 2017). In the current work, we conversely document a decrease in virally induced tonic PI3K drive, enhanced IgG class-switched cognate B-cell responses and NAb induction against a notably difficult-to-neutralize SIV. The exact hierarchy of binding between Nef and PI3K/mTORC2 components (including PI3K isoforms) and its alteration by Nef-G63E mutation remains to be investigated. Yet PI3Kp85 showed the strongest Nef-binding index in PLA assay (**Figure 4D**) presumably because of the SH3-binding PxxP domain in Nef which binds the SH3 domain of PI3Kp85 (Rickles et al., 1994). Decreased binding of Nef to its canonical target PI3Kp85 may precipitate into attenuated interactions of Nef with quintuple PI3K/mTORC2, resulting in fragile downstream Akt signaling. Such domino-triggering is also observed in APDS patients, from gain-of-function mutant p85 to p110 (Dornan et al., 2017), which similarly illustrates the fine-tuned nature of the PI3K/mTORC2/Akt signaling pathway.

A limitation of this study is the use of retrospective samples, posing constraints for detecting exact temporal changes in tonic viral B-cell perturbation. Related with this, it was not attainable to optimally time the sampling of lymph node cells in animals belonging to the subgroup of interest. Certain plasma and cellular samples were also unavailable in differing experiments. While partially addressed in *in vitro* reconstitution, much remains to be investigated for the observed polarity of Env-specific B-cell perturbation *in vivo*. Similarly, influence of the Nef-G63E phenotype on CD4^+^ T cells *in vivo* and its consequence on B-cell modulation remains largely elusive, and in the current work we focused mainly on molecular properties of the NAb-correlating Nef-G63E mutant strain. HIV/SIV infection uniquely shows a biphasic Tfh dysregulation of acute-phase destruction (Mattapallil et al., 2005; Moukambi et al., 2015; Moukambi et al., 2019) followed by dysregulated chronic-phase hyper-expansion (Lindqvist et al., 2012; Petrovas et al., 2012; Cubas et al., 2013; Xu et al., 2015). Further studies are therefore warranted to uncover how the Nef mutant, which is proapoptotic and displays ameliorated PI3K drive, influences Tfh responses at different phases of the infection. Gain-of-function mutations in PI3K-encoding genes appear to affect B cells more strongly (Asano et al., 2018). Nonetheless, it is noteworthy that they also evoke basal dysregulation in peripheral CD4^+^ T cells, which locally recapitulates dysregulation and death of HIV/SIV-infected CD4^+^ T cells, highlighting the importance of PI3K-mediated signaling in HIV/SIV infection. In any event, we surmise that specific animal models like the current one help to depict a temporal cascade of *in vivo* events leading to NAb induction, assisting human immunology. As an extension of this study extracting the NAb-involved molecular axis, manipulation/reconstitution experiments shall be designed.

In conclusion, we demonstrated in a non-human primate AIDS model that NAb induction against a difficult-to-neutralize SIV strain occurs after selection of a CD8^+^ T-cell escape variant with a reduced ability to drive excess PI3K/mTORC2 signaling. These results collectively offer an example of how NAb responses against immunodeficiency viruses can be shaped by both wings of adaptive immune pressure. Given the key role for Nef-mediated perturbation of PI3K/mTORC2 in NAb resistance, immune cell-intrinsic PI3K/mTORC2 manipulation may offer a new possibility to harness antiviral NAb responses. As human IEIs are increasingly becoming discovered with various immune-perturbing phenotypes and inheritance patterns (Jauch et al., 2023), search of other types of analogies between viral immune dysregulations and human IEIs may fuel the discovery of novel targets to modulate and harness immune responses in translational settings.

## Acknowledgments

This work was supported by Japanese Initiative for Progress of Research on Infectious Diseases for Global Epidemic (J-PRIDE) (JP19fm0208017 to H.Y.), Japan Program for Infectious Diseases Research and Infrastructure (Interdisciplinary Cutting-edge Research) (JP22wm0325006 to H.Y.), Research Programs on HIV/AIDS, Emerging and Re-emerging Infectious Diseases, and Global Health Issues (JP20fk0410022 and JP24fk0410066 to H.Y.; JP18fk0410003, JP20fk0410011, JP20fk0108125, JP20jm0110012, JP21fk0410035, and JP21jk0210002 to T.M.) from Japan Agency for Medical Research and Development, the Ministry of Education, Culture, Sports, Science, and Technology in Japan ([JSPS] KAKENHI Grants 17H02185, 18K07157 and 24K21287 to H.Y.; 21H02745 to T.M.), the Takeda Science Foundation (to H.Y.), the Imai Memorial Trust for AIDS Research (to H.Y.), and the Mitsui-Sumitomo Insurance Welfare Foundation (to H.Y.). We thank Fumiko Ono, Koji Hanari, Katsuhiko Komatsuzaki, Akio Hiyaoka, Hiromi Ogawa, Keiko Oto, Hirofumi Akari, Yasuhiro Yasutomi, Hiromi Sakawaki, Tomoyuki Miura and Yoshio Koyanagi for animal experiment assistance; Hiroyuki Kogure, Sayuri Seki and Julia R. Hirsiger for imaging cytometry technical assistance; Masako Nishizawa, Trang Thi Thu Hau, Shigeyoshi Harada, Takushi Nomura, Akiko Takeda, Taku Nakane, Nami Iwamoto, Taeko K. Naruse, Akinori Kimura and Mark S. de Souza for their help. H.Y. thanks Shoi Shi, Bettina Stolp, Jens V. Stein, Makoto Yamagishi and Yusuke Yanagi for conceptual discussions, and Jun Abe and Mike Recher for collaborative support.

## Author Contributions

H.Y. conceived/designed/performed study and wrote manuscript; T.M. provided animal cohort, co-edited manuscript and co-directed study.

## CRediT taxonomy

Conceptualization: HY

Methodology: HY

Validation: HY

Formal analysis: HY

Investigation: HY

Resources: HY, TM

Data curation: HY

Writing—original draft: HY

Writing—review & editing: HY, TM

Visualization: HY

Supervision: HY, TM

Project administration: HY

Funding acquisition: HY, TM

## Materials and Methods

### Material availability

This study did not generate new unique reagents.

### Data and code availability

Data were analyzed using existing computational packages. The accession number for the transcriptome data [WT SIV_mac239_-infected HSC-F cell line (*n* = 3), Nef-G63E SIV_mac239_-infected HSC-F cell line (*n* = 3) and uninfected HSC-F cell line (*n* = 3)] has been deposited under the accession number GenBank: GSE65806.

### Rhesus macaques retrospectively utilized for samples

Seventy Burmese rhesus macaques (*macaca mulatta*) (57 males and 13 females) were retrospectively analyzed in this study. Experiments were previously carried out (Matano et al., 2004; Yamamoto et al., 2007; Iseda et al., 2016; Nomura et al., 2012; Ishii et al., 2012; Takahashi et al., 2013; Nakane et al., 2013; Shi et al., 2013; Iwamoto et al., 2014; Terahara et al., 2014) in the Tsukuba Primate Research Center, National Institutes of Biomedical Innovation, Health and Nutrition (NIBIOHN) with the help of the Corporation for Production and Research of Laboratory Primates and the Institute for Frontier Life and Medical Sciences, Kyoto University (IFLMS-KU) after approval by the Committee on the Ethics of Animal Experiments of NIBIOHN and IFLMS-KU under the guidelines for animal experiments at NIBIOHN, IFLMS-KU and National Institute of Infectious Diseases in accordance with the Guidelines for Proper Conduct of Animal Experiments established by Science Council of Japan (http://www.scj.go.jp/ja/info/kohyo/pdf/kohyo-20-k16-2e.pdf). The experiments were in accordance with the “Weatherall report for the use of non-human primates in research” recommendations (https://royalsociety.org/topics-policy/publications/2006/weatherall-report/). Animals were housed in adjoining individual primate cages allowing them to make sight and sound contact with one another for social interactions, where the temperature was kept at 25°C with light for 12 hours per day. Animals were fed with apples and commercial monkey diet (Type CMK-2, Clea Japan, Inc.). Blood collection and virus challenge were performed under ketamine anesthesia.

### Cells and viruses

MT4-R5 cells were maintained in RPMI1640 (Invitrogen) supplemented with 10% fetal bovine serum (FBS) (Clontech) and antibiotics. HSC-F cells (cynomolgus CD4^+^ T-cell line) and HSR5.4 cells (rhesus macaque CD4^+^ T-cell line) were maintained in RPMI1640 (Invitrogen) supplemented with 10% fetal bovine serum (FBS) (Clontech), human interleukin-2 (IL-2) (10 IU/ml, Roche), 4-(2-hydroxyethyl)-1-piperazineethanesulfonic acid (HEPES) (Invitrogen) and 2-mercaptoethanol (Gibco). SIV_mac239_ molecular clone derivative was generated by mutagenesis PCR (Agilent Technologies) using the primers listed. For virus preparation, COS-1 cells were transfected with pBR_mac239_ proviral DNA using FuGENE 6 (Promega). At 48 h post-transfection, culture supernatants were harvested, centrifuged and filtered through a 0.45-μm pore-size filter (Merck Millipore). To titrate infectivity, prepared viruses were serially diluted and infected on HSC-F cells in 96-well plates (Falcon) in quadruplicate. At 10 days post-infection, the endpoint was determined using SIV p27 antigen enzyme-linked immunosorbent assay (ELISA) kit (ABL), and virus infectivity was calculated as the 50 percent tissue culture infective dose (TCID_50_) according to the Reed-Muench method.

### Identification of SIV_mac239_-NAb inducers

Burmese rhesus macaques previously challenged with the highly pathogenic molecular clone virus SIV_mac239_ (*n* = 70) were retrospectively examined approximately up to 2 years for their plasma NAb profiles. Animals were challenged intravenously with 1,000 TCID_50_ of SIV_mac239_. In the current study, virological and immunological profiles were compared between the newly identified NAb inducers (*n* = 9) and representative NAb non-inducers (*n* = 19). Sex distribution of the NAb inducers (eight males and one female) did not significantly differ with the total non-inducers (49 males and 12 females, *P* = 0.99 by Fisher’s exact test). These NAb non-inducers and eight NAb inducers except R06-032 were previously partially reported for their plasma viral loads (Yamamoto et al., 2007; Iseda et al., 2016; Nomura et al., 2012; Takahashi et al., 2013; Nakane et al., 2013; Iwamoto et al., 2014) (**Figure 1B**). MHC-I haplotypes and alleles were determined by reference strand-mediated conformation analysis, PCR-SSP (PCR amplification utilizing sequence-specific priming) and cloning as described (Naruse et al., 2010). MHC-I binding prediction was made on April 27, 2013 using the IEDB analysis resource NetMHCpan tool (Hoof et al., 2009). Alleles of interest in the study have been previously identified in macaques (Nomura et al., 2012; Evans et al., 1999; Sette et al., 2012). Macaques R06-032, R03-015, R01-010 and R05-010 received a prime-boost vaccination (Matano et al., 2004) composed of a DNA prime/intranasal Sendai virus vector expressing SIV_mac239_ Gag (SeV-Gag). R03-015, R06-019, R06-038 and R10-001 received 300 mg of non-specific rhesus IgG at day 7 post-SIV_mac239_ challenge as an experimental control in our previous reports (Yamamoto et al., 2007; Iseda et al., 2016; Nakane et al., 2013).

### Plasma viral load quantitation

Plasma viral RNA samples were extracted with High Pure Viral RNA kit (Roche Diagnostics). Serial fivefold sample dilutions were amplified in quadruplicate by reverse transcription and nested PCR using SIV_mac239_ *gag*-specific primers to determine end point via the Reed-Muench method as described previously (Matano et al., 2004; Iseda et al., 2016). The lower limit of detection is approximately 400 viral RNA copies/ml plasma. Viral loads have been previously partially reported (Yamamoto et al., 2007; Nomura et al., 2012; Takahashi et al., 2013; Nakane et al., 2013; Iwamoto et al., 2014).

### SIV_mac239_-specific neutralization assay

NAbs were titrated as described (Yamamoto et al., 2007; Iseda et al., 2016). Serial twofold dilutions of heat-inactivated plasma, or polyclonal IgG affinity-purified with Protein G Sepharose 4 Fast Flow (GE Healthcare) from heat-inactivated and filtered plasma were mixed with 10 TCID_50_ of SIV_mac239_ at a 1:1 ratio (5 μl:5 μl) in quadruplicate. After 45-min incubation at room temperature, the 10 μl mixtures were added into 5 x 10^4^ MT4-R5 cells/well in 96-well plates. Progeny virus production in day 12 culture supernatants was examined by SIV p27 ELISA (ABL) to determine 100% neutralizing endpoint. The lower limit of titration is 1:2. Neutralization in three out of four wells at a dilution of 1:2 is defined as marginally NAb-positive (< 1:2). Results were comparable when the same assay was performed with macaque HSC-F cells (Akari et al., 1996) as targets. NAb inducers R01-012, R02-004, R01-010, R02-007 and NAb non-inducer R01-011 were previously partially reported for their NAb titers measured with the same method using MT4 cells as targets (Kawada et al., 2007), which derived comparable results. For assessment of neutralizing activity in IgG, SIV_mac239_-specific IgGs purified from pools of plasma with SIV_mac239_-specific NAb titers were obtained from each animal as described (Yamamoto et al., 2007). After complement heat-inactivation at 56°C, 30 min and 0.45 μm filtration, IgGs were purified by Protein G Sepharose 4 Fast Flow (GE Healthcare) and concentrated by Amicon Ultra 4, MW 50,000 (Millipore) to 30 mg/ml and similarly examined for their 10 TCID_50_ SIV_mac239_ killing titers on MT4-R5 cells.

### SIV Env-specific IgG ELISA and immunoblotting

Plasma Env-specific IgG titers were measured as described (Nakane et al., 2013). SIV_mac251_ Env gp120 (ImmunoDiagnostics) were coated on 96-well assay plates (BD) at 1000 ng/ml (100 μl/well). Wells were prewashed with phosphate-buffered saline (PBS), blocked with 0.5% bovine serum albumin (BSA)/PBS overnight and plasma samples were incubated at a 1:20 dilution (5 μl:95 μl) for 2 hours. Wells were washed with PBS and SIV Env-bound antibodies were detected with a horseradish peroxidase (HRP)-conjugated goat anti-monkey IgG (H+L) (Bethyl Laboratory) and SureBlue TMB 1-Component Microwell Peroxidase Substrate (KPL). Absorbance at 450 nm was measured. Samples from week 0 pre-challenge and month 3, month 6 and year 1 post-challenge were assessed in duplicate. Week 0 average values were subtracted from corresponding later time point values for calibration. For immunoblotting, SIV virion-specific IgGs in plasma were detected with a SIV_mac239_-cross-reactive western blotting system (ZeptoMetrix). In the NAb non-inducers, samples from those close to rapid progression (euthanized due to AIDS progression within approximately 1 year) known for low plasma anti-SIV reactivity (Hirsch et al., 2004; Nakane et al., 2013) were not included.

### Sequencing

Sequencing was performed as described (Matano et al., 2004; Iseda et al., 2016). Viral cDNA fragments spanning from nt (nucleotide) 4829 to nt 7000, nt 6843 to nt 8831 and nt 8677 to nt 10196 in SIV_mac239_ (GenBank accession number MM33262) covering SIV *env* and *nef* were amplified from plasma viral RNA by nested RT-PCR using Prime-Script one-step RT-PCR kit v2 (TaKaRa) and KOD-Plus v2 (Toyobo). PCR products were either directly sequenced or subcloned with a TOP10-transforming TOPO blunt-end cloning system (Invitrogen). Sequencing was performed using dye terminator chemistry with an ABI 3730 DNA sequencer (Applied Biosystems). On average, 15 clones were obtained per sample and 20 clones were assessed when *nef* mutations of interest in early time points were subdominant.

### SIV_mac239_-specific CD8^+^ T-cell responses

Virus-specific CD8^+^ T-cell frequencies were measured as described (Matano et al., 2004; Iseda et al., 2016). Peripheral blood mononuclear cells (PBMCs) were cocultured for 6 hours with autologous herpesvirus papio-immortalized B lymphoblastoid cell lines (B-LCLs) pulsed with Nef peptides (Sigma-Aldrich Japan) at 1 μM concentration or as indicated otherwise under GolgiStop (monensin, BD) presence. Intracellular gamma interferon (IFN-γ) staining was performed using Cytofix/Cytoperm kit (BD) and the following conjugated anti-human monoclonal antibodies (mAbs): anti-CD4-FITC (M-T477, BD Pharmingen), anti-CD8-PerCP (SK1, BD Biosciences), anti-CD3-APC (SP34-2, BD Pharmingen) and anti-IFN-γ-PE (4S.B3, BioLegend). Specific CD8^+^ T-cell frequencies were determined by subtracting nonspecific IFN-γ^+^ CD8^+^ T-cell frequencies from those after peptide-specific stimulation; frequencies beneath 100 cells/million PBMCs were considered negative. Cells acquired by FACS Canto II (BD) were analyzed by FACS Diva (BD) and FlowJo (Treestar). Approximately 1 x 10^5^ PBMCs were gated per test.

### Nef-mediated signaling perturbation analysis

Virus supernatants obtained from COS-1 cells after transfection with WT or mutant SIV_mac239_ molecular clones were used for infection of CD4^+^ T-cell lines, cynomolgus macaque-derived HSC-F and rhesus macaque-derived HSR5.4 (Akari et al., 1996). QuikChange II XL site-directed mutagenesis kit (Agilent Technologies) was used to construct mutant SIV_mac239_ molecular clones possessing *nef* mutations Nef-G63E (G-to-A mutation at nt 9520) and Nef-G63R (G-to-A mutation at nt 9519) from the WT SIV_mac239_ molecular clone (Kestler et al., 1991) (nt number from GenBank accession number M33262). Cells (1 x 10^5^ cells/well in U-bottomed 96-well culture plates [BD]) were infected with WT or mutant SIV_mac239_ cultured in RPMI-1640 medium supplemented with 10% fetal bovine serum (FBS) for indicated periods at MOI 0.1 (intracellular signaling analysis) or 0.001 (supernatant analysis). Culture supernatants were subjected to measurement of SIV capsid p27 concentrations by ELISA. Harvested cells were fixed and permeabilized with Cytofix/Cytoperm kit, washed twice and subjected to immunostaining. The following antibodies were used; anti-SIV_mac251_ Gag p27 mAb (ABL) manually conjugated with Alexa 647 (Life Technologies), anti-SIV_mac251_ Nef mAb (clone 17, epitope peptide corresponding to SIV_mac239_ Nef 71-80 not including the residue G63: Thermo Fisher Scientific/Pierce) manually conjugated with Alexa 488 or Alexa 647, anti-human CD3-APC-Cy7 (SP34-2, BD Pharmingen), anti-human CD4-PerCP (L200, BD Pharmingen), anti-human HLA-ABC-FITC (G46-2.6, BD Pharmingen), anti-human CXCR4-APC (12G5, BioLegend), BST2-Alexa 647 (RS38E, BioLegend), Alexa 488-conjugated anti-phospho-Akt (Ser473) (D9E, CST) and Alexa 488-conjugated anti-phospho-Akt (Thr308) (C31E5E, CST). Cells acquired by FACS Canto II were analyzed by FACS Diva and FlowJo. Approximately 5 x 10^4^ cells were gated for each test.

### Ferritin ELISA

Plasma ferritin levels in NAb inducers and control animals were analyzed by monkey ferritin sandwich ELISA kit (LS Bio) according to the manufacturer’s instructions.

### PI3K stimulation assay

5 x 10^4^ HSC-F cells were infected with WT or Nef-G63E mutant SIV_mac239_ at MOI 0.2 and cultured for 1 day in medium supplemented with 10% (normal), 1% (1/10 starvation) or 0.1% (1/100 starvation) FBS. For ligand stimulation (**Figure 4-figure supplement 1C**), cells at the end of 1-day culture were pulsed for 20 minutes with 40 ng/ml of recombinant human IFN-γ (Gibco/Thermo Fisher Scientific), 100 IU/ml of recombinant human IL-2 (Roche Diagnostics) or 10 μg/ml of SIV_mac251_ Env gp130 (ImmunoDx). Cells were intracellularly stained using Cytofix/Cytoperm kit with anti-SIV_mac251_ Nef mAb or anti-SIV_mac251_ p27 mAb manually conjugated to Alexa 647 and PE-conjugated anti-human phospho-Akt (Ser473) (D9E, CST) or Alexa 488-conjugated anti-human phospho-Akt (Ser473). Cells acquired by FACS Canto II were analyzed by FACS Diva and FlowJo. Approximately 7 x 10^4^ HSC-F cells were gated per test.

### Transcriptome analysis

Total RNAs were extracted using RNeasy Plus Mini kit (Qiagen) from 2 x 10^6^ HSC-F cells 1 day after infection with WT or Nef-G63E mutant SIV_mac239_ at MOI 5. Negative control RNA samples were extracted from 2 x 10^6^ uninfected HSC-F cells after culture with the same condition. Three sets of experiments were performed. Total RNA samples were subjected to a quality control (QC) analysis using an Agilent 2100 Bioanalyzer. The obtained amounts of total RNAs were: 12.37 ± 0.39 (uninfected), 8.80 ± 0.44 (WT) and 9.02 ± 0.17 (Nef-G63E) μg (*P* = 0.66 for WT versus Nef-G63E by unpaired t test). In all samples, two bands of 18S and 28S rRNA were confirmed and the RNA integrity number (RIN) was 10. 500 ng of total RNA samples were processed with GeneChip WT Plus reagent (Affymetrix/Thermo Fisher Scientific) to produce 150 μl of fragmented and labeled cDNA samples. These were incubated with a Human Gene 2.0 ST Array (Affymetrix/Thermo Fisher Scientific) for 16 hours, 60 rpm at 45°C using GeneChip hybridization oven 645 (Affymetrix/Thermo Fisher Scientific). Results were scanned with GeneChip Scanner 3000 7G (Affymetrix/Thermo Fisher Scientific) and processed with Affymetrix Expression Console Software (Affymetrix/Thermo Fisher Scientific) according to the manufacturer’s instructions. Expression values were normalized by the RMA method. Genes above mean background expression within the cognate sample and showing significant difference between WT and Nef-G63E SIV (*P* < 0.05 via unpaired t test for log_2_-transformed values, 768 candidates) were determined. Akt-related genes exhibiting a change of approximately 10% or more were representatively extracted and manually aligned by the authors.

### Peripheral CD4^+^ T-cell surface staining

Cryopreserved/thawed PBMCs were stained for 30 minutes at 4°C with the following reagents or conjugated anti-human mAbs: Live/Dead Aqua (Life Technologies), anti-CD4-PerCP (L200, BD Pharmingen), anti-CD8-APC-Cy7 (RPA-T8, BD Pharmingen), anti-CD3-Alexa 700 (SP34-2, BD Pharmingen), anti-CD95-PE-Cy7 (DX2, eBioscience), anti-CXCR5-PE (87.1, eBioscience), anti-PD-1-Brilliant Violet 421 (EH12.2H7, BioLegend) and anti-CXCR3-Alexa 488 (G025H7, BioLegend). Cells acquired by FACS LSRII Fortessa (BD) were analyzed by FACS Diva and FlowJo. Approximately 1.5 x 10^5^ PBMCs were gated per test.

### Proximity ligation assay (PLA)

A flow cytometry-based arrangement of proximity ligation assay (PLA) (Avin et al., 2017) was performed to quantitatively assess Nef binding to candidate interacting molecules. 1 x 10^5^ HSC-F cells were infected with SIV_mac239_ at MOI 0.05 in U-bottomed plates and permeated with Cytofix/Perm kit (BD Biosciences). After two washes, they were resuspended in 0.5% BSA/PBS for prevention of experimental procedure-related loss and stained with mouse anti-SIV_mac251_ Nef mAb (clone 17, Thermo Fisher Scientific/Pierce) or mouse IgG1 isotype control mAb (P3.6.2.8.1, Abcam) in combination with either of the following rabbit antibodies: anti-GβL (86B8, CST), anti-mTOR (7C10, CST), polyclonal anti-human PI3K p85 (Merck Millipore), anti-PI3 Kinase p110α (C73F8, CST) or rabbit IgG1 isotype control mAb (DA1E, CST). Antibody-stained cells were subsequently probed with Duolink In Situ PLA Probe anti-mouse PLUS and anti-mouse MINUS probes (Sigma/Merck). Next, they were detected for intermolecular binding using Duolink flow PLA detection kit (Green) (Sigma/Merck) with a reaction time of 100 minutes for post-mouse/rabbit probe linking amplification. Finally, these PLA-subjected cells were additionally stained with anti-SIV_mac251_ Gag p27 mAb manually conjugated with Alexa 647 and anti-CD4-PerCP for 30 minutes for 20 minutes at 4°C. Cells acquired by FACS Canto II were analyzed by FACS Diva and FlowJo. Approximately 1 x 10^5^ cells were gated per test. Binding index (Y axis) was calculated by deriving the deviation from the summation of: {baseline P: (MFI of background reaction with mouse isotype control/rabbit isotype control) + anti-Nef antibody-derived background Q: [(MFI of reaction with mouse anti-Nef/rabbit isotype control) - (MFI of background reaction with mouse isotype control/rabbit isotype control)] + anti-binding partner antibody-derived background R:[(MFI of reaction with mouse isotype control/rabbit anti-binding partner molecule) - (MFI of background reaction with mouse isotype control/rabbit isotype control)]}.

### Co-immunoprecipitation analysis

1 x 10^5^ HSC-F cells were infected with SIV_mac239_ at MOI 0.05 in U-bottomed plates in quadruplicate for each SIV strain for three days and pooled for each for acquiring cell pellets. These pellets were lysed with Capturem IP & Co-IP Kit lysis buffer (Takara) and portions for each were immunoprecipitated by anti-Sin1 mAb (D7G1A, CST) (1:50 dilution, 20 minutes, room temperature). Spin column membrane-bound immunoprecipitates were obtained by centrifugation with Capturem IP & Co-IP Kit (Clontech/Takara). Whole cell lysates and Sin1-immunoprecipitated products were subjected to SDS-polyacrylamide gel electrophoresis separation on a Mini Protean TGX 4-15% gel (BioRad) and transferred to a Immun-Blot PVDF membrane (BioRad). Immunoblotting was performed by mouse anti-SIV_mac251_ Nef mAb (clone 17, Thermo Fisher Scientific/Pierce) primary antibody probing (1:1000 dilution, 18 hours, 4°C) and Mouse TrueBlot ULTRA anti-mouse Ig HRP (eB144, Rockland Immunochemicals) secondary antibody incubation (1:1000 dilution, 30 minutes, room temperature). Nef-specific bands in infected controls were visualized by enhanced chemiluminescence using SuperSignal West Pico PLUS (Thermo Fisher Scientific). Bands auto-detectable by Image Lab software (BioRad) were analyzed for signal intensities. Uncropped images are provided as supplementary data.

### Cell death assay

Infected cell apoptosis was measured by methods previously described (Jauch et al., 2023). 1 x 10^5^ HSC-F cells were infected with SIV_mac239_ at MOI 0.1 in U-bottomed plates (six wells/control) for each SIV strain for three days. These cells were stained with CD4-FITC (M-T477, BD Pharmingen) for 15 minutes at room temperature, washed once and next stained with APC Annexin V (Biolegend) in Annexin V Binding Buffer (Biolegend) for 20 minutes at room temperature. Stained reactions were then diluted fivefold with Annexin V Binding Buffer and subjected to analysis. Cells acquired by FACS Lyric were analyzed by FACS Suite and FlowJo. Approximately 1 x 10^4^ HSC-F cells were gated per test.

### Quantitative *in vivo* imaging flow cytometry

Cryopreserved/thawed lymph node cells (LNCs) from several persistently SIV-infected macaques used in previous experiments (Nakane et al., 2013; Takahashi et al., 2013; Iwamoto et al., 2014) were seeded in V-bottomed 96-well plates (Nunc) and blocked with 25 μg/ml of anti-human CD4 (clone L200: BD) in 100 μl volume for 15 minutes at 4°C. After three washes, they were stained with 10 μg/ml of recombinant SIV_mac239_ Env gp140-biotin (Immune Technology) for 30 minutes at 4°C. Cells were then stained with anti-CD19-PC5.5 (J3-119, Beckman Coulter), Live/Dead Aqua (Invitrogen) and streptavidin-APC-Cy7 (BioLegend) for 30 minutes at 4°C. After two washes, cells were processed with Cytofix/Cytoperm kit (BD) for 20 minutes at 4°C, washed twice, and next intracellularly stained with PE-conjugated anti-human phospho-Akt (Ser473) (D9E, CST) and anti-SIV_mac251_ Nef mAb (clone 17, Thermo Fisher Scientific/Pierce) manually conjugated with Alexa 488 or mouse IgG1 isotype control mAb (P3.6.2.8.1, Abcam) for 30 minutes at 4°C. After final two washes, cells were suspended in 0.8% PFA/PBS. All centrifugations for washing (1,200 x g, 2 minutes) were performed at 4°C. Cells were subjected to image acquisition with Image Stream X MKII imaging flow cytometer (Amnis/Merck Millipore/Luminex) and analyzed with IDEAS 6.3 software (Amnis/Merck Millipore/Luminex). Approximately 5 x 10^5^ LNCs were acquired for analyses (**Figure 5**). A custom-implemented linear discriminant analysis-based machine learning module (Luminex) was utilized for verifying candidate Nef staining signal-related secondary parameters (**Figure 5** and **Figure 5-figure supplement 1A**) for their efficacy of target population separation.

### B-cell Nef invasion *in vitro* assay

For Nef invasion assay, 5 x 10^4^ HSC-F cells were infected at MOI 1 for 48 hours. These were then additionally cocultured with 7.5 x 10^4^ Ramos B cells for 12 hours with or without addition of macaque T cell-stimulating anti-CD3 antibody (FN-18, Abcam) at 1 μg/ml. Cells were double-stained with Alexa 488-conjugated mouse anti-Nef (clone 17: Thermo Scientific/Pierce) and anti-CD19-PC-5.5 (J3-119, Beckman Coulter). For graphical visualization (**Figure 6A**), interacting Nef^+^ infected cell-invaded B cell doublet images acquired with ImageStream X MKII (Amnis/Merck Millipore) were analyzed by IDEAS 6.3 (Amnis/Merck Millipore/Luminex) with the defined noise-cancelling gating (via gating Nef pixel signal contrast^lo^-Nef mean pixel per object^hi^ cells). For Nef invasion modulation verification on conventional flow cytometry, the aforementioned culture condition was similarly used. Anti-CD3 antibody (FN-18, Abcam) addition at 1 μg/ml and phorbol 12-myristate 13-acetate (PMA, Sigma) plus ionomycin (Sigma) addition (10 ng/ml and 50 ng/ml, respectively) l were evaluated. Cells were stained with Alexa 647-conjugated mouse anti-Nef (clone 17: Thermo Scientific/Pierce), anti-CD19-PC-5.5 (J3-119, Beckman Coulter), anti-human phospho-Akt (Ser473)-Alexa 488 (D9E, CST) and Live/Dead Near-IR (Invitrogen) for dead cell exclusion. Dead cell frequencies were less than 3%. For viral phenotype evaluation, 1 x 10^5^ HSC-F cells (surface CD40L^+^) were pre-infected at MOI 0.2 with WT or Nef-G63E mutant SIV_mac239_ six hours before coculture. 1.5 x 10^5^ CD20^+^ B cells, positively selected from fresh PBMCs of uninfected rhesus macaques via anti-CD20 microbeads (Miltenyi Biotec), were cocultured with these HSC-F cells at an E:T ratio of 2:3 for 3 days in the presence of 40 ng/ml carrier-free recombinant human IL-4 (R&D Systems). Cells were surface-stained with anti-human CD20-PerCP (2H7, BioLegend) and intracellularly stained using Cytofix/Cytoperm kit with anti-SIV_mac251_ Nef mAb manually conjugated to Alexa 488 and Alexa 647-conjugated anti-human phospho-Akt (Ser473) (D9E, CST). For **Figure 6B** and **Figure 6C**, Cells acquired by FACS Canto II were analyzed by FACS Diva and FlowJo. Approximately 5 x 10^4^ CD20^+^ primary B cells and 3 x 10^4^ CD19^+^ Ramos B cells were analyzed.

### Peripheral SIV Env-specific B-cell responses

Peripheral SIV Env gp140-specific memory B cells (B_mem_) and plasmablasts (PBs) were measured by flow cytometry with procedures modified from previous reports and our experience (Silveira et al., 2015; Hau et al., 2022). First, cryopreserved/thawed PBMCs seeded in V-bottomed 96-well plates (Nunc) were blocked with 25 μg/ml of anti-human CD4 (clone L200: BD) in 100 μl volume for 15 minutes at 4°C. This concentration attains blockade of promiscuous SIV Env binding to CD4 comparable to levels by magnetic depletion of CD3^+^ T cells (data not shown). After three washes, they were next stained with 10 μg/ml of recombinant SIV_mac239_ Env gp140-biotin (Immune Technology) for 30 minutes at 4°C. They were subsequently stained with the following anti-human antibodies and fluorochromes for 30 minutes at 4°C with combinations as follows; PB/B_mem_: anti-CD3-APC-Cy7 (SP34-2, BD Pharmingen), anti-CD8-APC-Cy7 (RPA-T8, BD Pharmingen), anti-CD14-APC-Cy7 (M5E2, BioLegend), anti-CD16-APC-Cy7 (3G8, BioLegend), streptavidin-Brilliant Violet 421 (BioLegend), anti-CD19-PC5.5 (J3-119, Beckman Coulter) and Live/Dead Aqua (Invitrogen); PB: anti-CD10-APC-Cy7 (HI10a, BioLegend), anti-CD11c-PE-Cy7 (3.9, BioLegend), anti-CD123-PE-Cy7 (6H6, BioLegend) and anti-HLA-DR-PE-Texas Red (TU36, Invitrogen); B_mem_: anti-CD27-PE/Dazzle 594 (M-T271, BioLegend), anti-CD10-PE-Cy7 (HI10a, BioLegend), anti-IgD-FITC (DaKo), anti-CD38-Alexa 647 (AT1, Santa Cruz Biotechnology), anti-IgG-Alexa 700 (G18-145, BD Pharmingen) and anti-CD138-PE (DL-101, eBioscience). After two washes, B_mem_ samples were suspended in 0.8% PFA/PBS. For PB staining, surface-stained cells were further processed with Cytofix/Cytoperm kit (BD) for 20 minutes at 4°C, washed twice, and next intracellularly stained with 10 μg/ml of recombinant SIV_mac239_ Env gp140-biotin (Immune Technology) for 30 minutes at 4°C. After three washes, they were next stained with anti-human IgG-PE (G18-145, BD), anti-human/mouse IRF4-eFluor 660 (3E4, eBioscience), anti-human Ki67-Alexa 700 (B56, BD Pharmingen) and streptavidin-Alexa 488 (BioLegend) for 30 minutes at 4°C. After final two washes, cells were suspended in 0.8% PFA/PBS. Cells acquired with FACS LSRII Fortessa were analyzed with FACS Diva and FlowJo. Approximately 1.5 x 10^5^ PBMCs were acquired for B_mem_ and 4 x 10^5^ PBMCs were acquired for PB analyses (**Figure 6**). All centrifugations for washing (1,200 x g, 2 minutes) were performed at 4°C. Env-specific PB frequencies were 0 cells/million PBMCs for pre-challenge samples in all animals examined, ruling out background confounding for high-sensitivity quantitation of the population.

### Statistical Analysis

Analyses were performed via Prism 8 (GraphPad Software). *P* < 0.05 were considered significant in two-tailed unpaired t tests, paired t tests, Mann Whitney U tests, Fisher’s exact tests, Wilcoxon signed-rank tests, one-way ANOVA with Tukey’s post-hoc multiple comparison tests and two-way ANOVA with Sidak’s post-hoc multiple comparison tests. For analysis of Nef-invaded B-cell pAkt levels, median fluorescence intensities (mFIs) were analyzed due to relatively small numbers of Nef^+^ B cells. Analyses involving pAkt Ser473 levels all derived comparable results between mFI and MFI for WT versus Nef-G63E virus. Machine learning-based rating of Nef signal parameters was verified with IDEAS 6.3 Machine Learning Module (Luminex).

## Supplemental Information Legends

**Fig. 1-Supplement 1.**
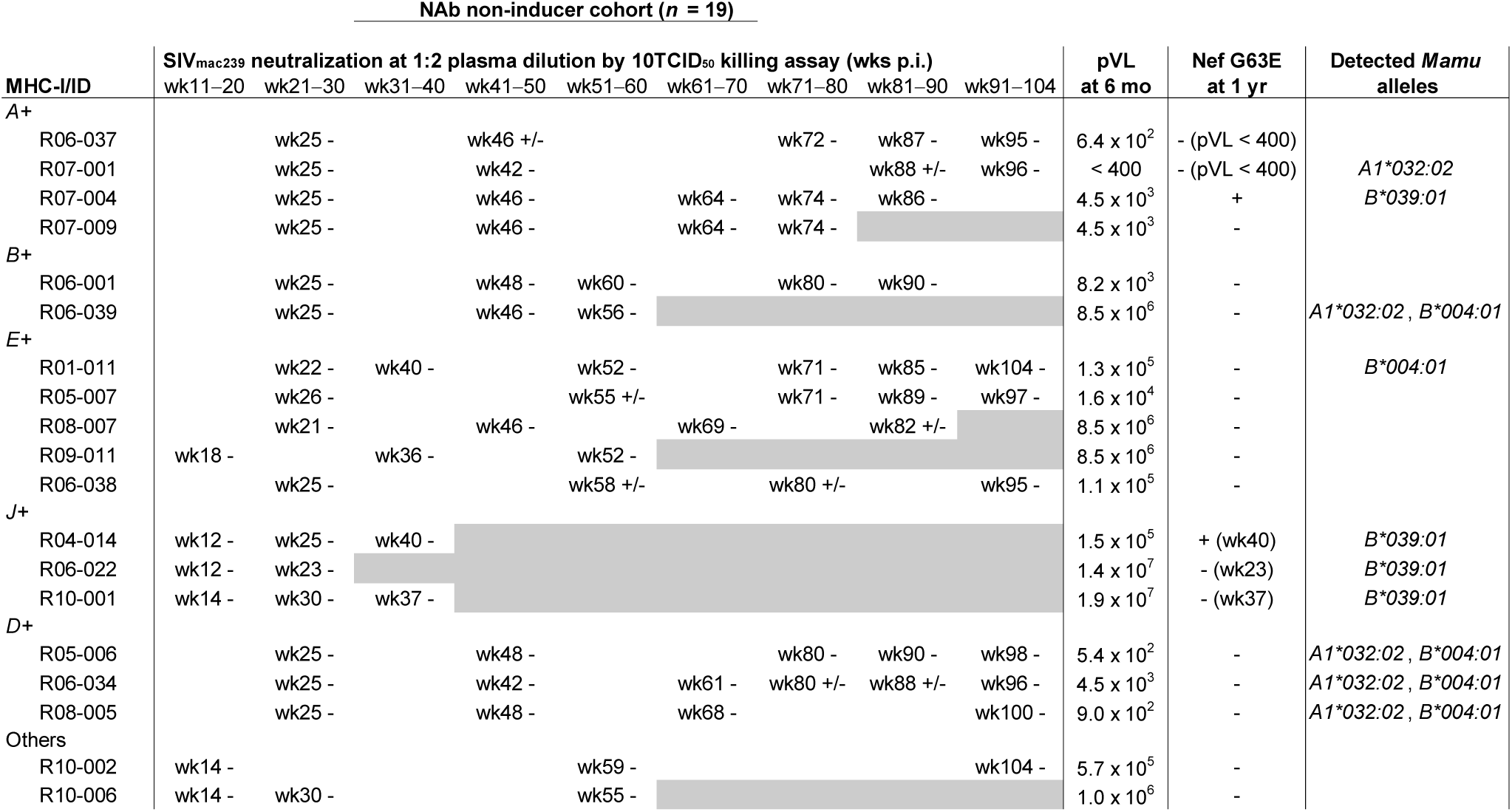
Profile of SIV_mac239_-NAb non-inducer subgroup cohort. Summary of a SIV_mac239_-NAb non-inducer subgroup cohort (*n* = 19), composed from previously partially characterized, MHC class I haplotype-balanced naïve animals. Symbols A+, B+, E+, J+ and D+ represent possession of MHC-I haplotypes *90-120-Ia*, *90-120-Ib*, *90-010-Ie*, *90-088-Ij* and *90-010-Id*, respectively. For the titers, - represents negative; +/- represents neutralization in three out of four wells in a quadruplicate test. Gray shadings indicate euthanasia due to AIDS progression. Mamu-A and Mamu-B alleles of interest in this study are listed. These NAb non-inducers were previously partially reported for their plasma viral loads and MHC-I alleles (Nomura et al., 2012; Takahashi et al., 2013; Nakane et al., 2013; Iwamoto et al., 2014). R01-011 was previously partially reported for NAb titers measured with the same method using MT4 cells as targets (Kawada et al., 2007), which derived comparable results.

**Fig. 1-Supplement 2.**
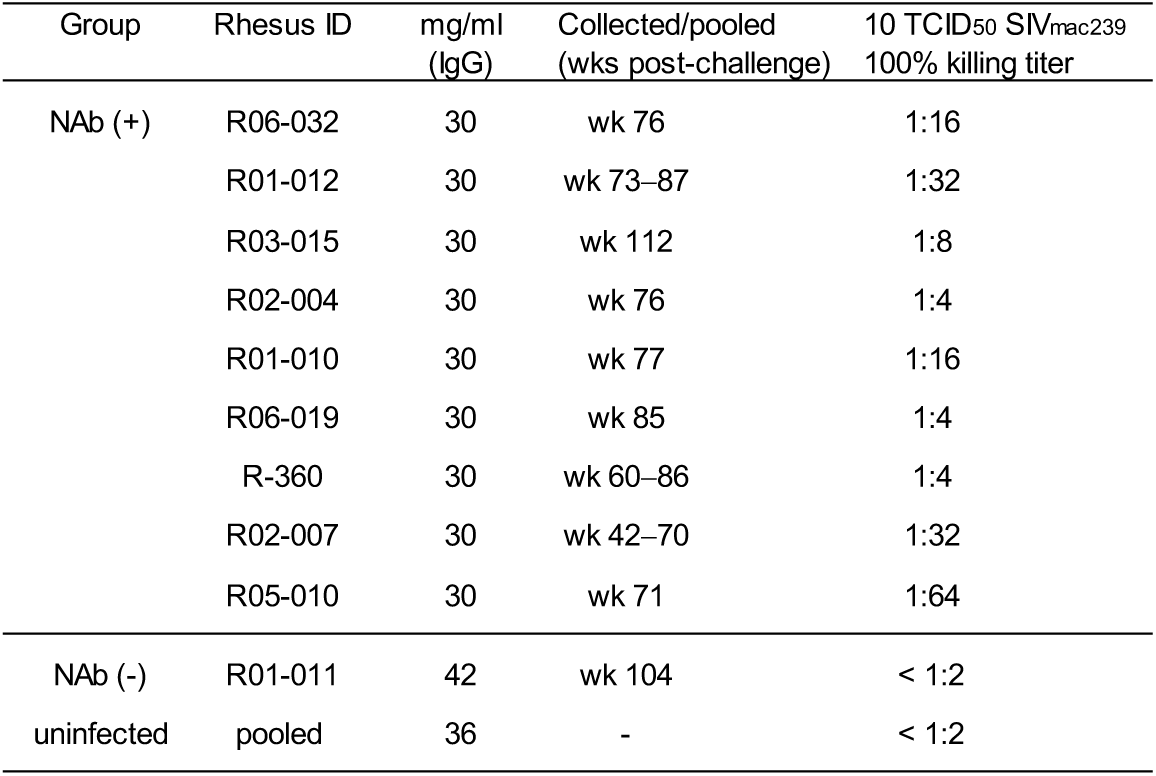
IgG class-switched NAbs in the inducers. Titration of SIV_mac239_-neutralizing IgGs in the NAb inducers. SIV_mac239_-specific IgGs, purified from pools of plasma with SIV_mac239_-specific NAb titers, were obtained from each animal as described (Yamamoto et al., 2007) and examined for their neutralizing activity by 10 TCID_50_ SIV_mac239_ killing on MT4-R5 cells. NAb-negative anti-SIV IgG and control IgG were prepared from a representative animal (R01-011) with NAb-negative plasma and pooled plasma of uninfected rhesus macaques, respectively.

**Fig.1-Supplement 3.**
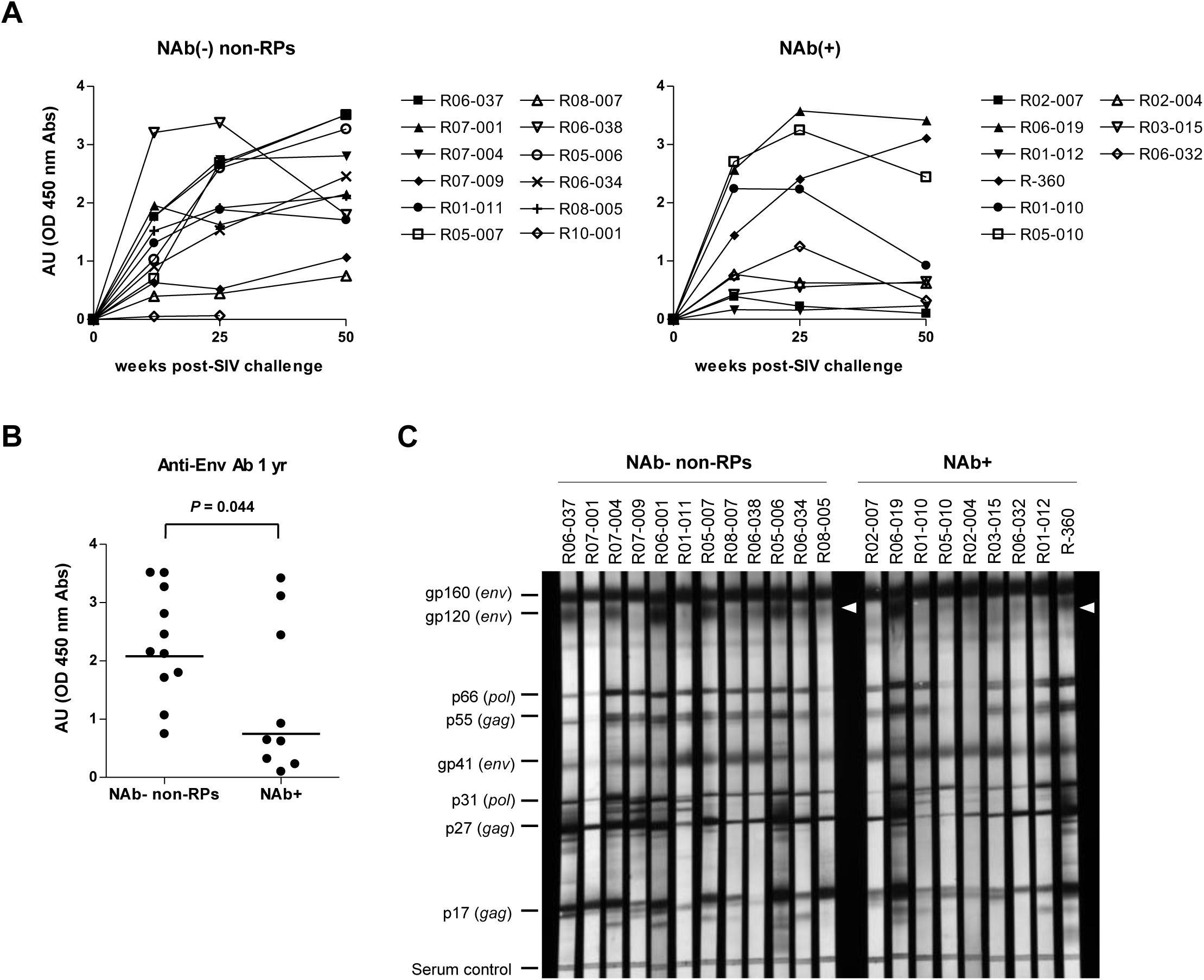
Anti-SIV binding IgG profiles in NAb inducers and non-inducers. **(A)** Temporal changes in SIV_mac251_ Env gp120-specific antibody titers in the NAb non-inducers without rapid progression [NAb-negative non-rapid progressors, NAb(-) non-RPs] (left, *n* = 11) and NAb inducers [NAb(+)] (right, *n* = 9). NAb non-inducers showing rapid progression (euthanized with AIDS onset within approximately 1 year) were not included except for macaque R10-001 (shown in the left for aligned comparison). Samples for non-RPs R06-001 and R10-002 were unavailable. **(B)** Unpaired t test of log-transformed anti-SIV Env Ab titers in NAb(-) non-RPs and NAb inducers. Bars represent geometric mean absorbance within each group. **(C)** Lysed SIV_mac_ virion linear antigen-binding at year 1 p.i. in NAb(-) non-RPs (left, *n* = 12) and NAb inducers (right, *n* = 9). Plasma anti-SIV_mac251_ IgGs were detected using a commercial western blotting system against the parental strain SIV_mac251_ (ZeptoMetrix). White arrowheads indicate Env gp120-specific bands. Experiments were performed twice with comparable results.

**Fig. 1-Supplement 4.**
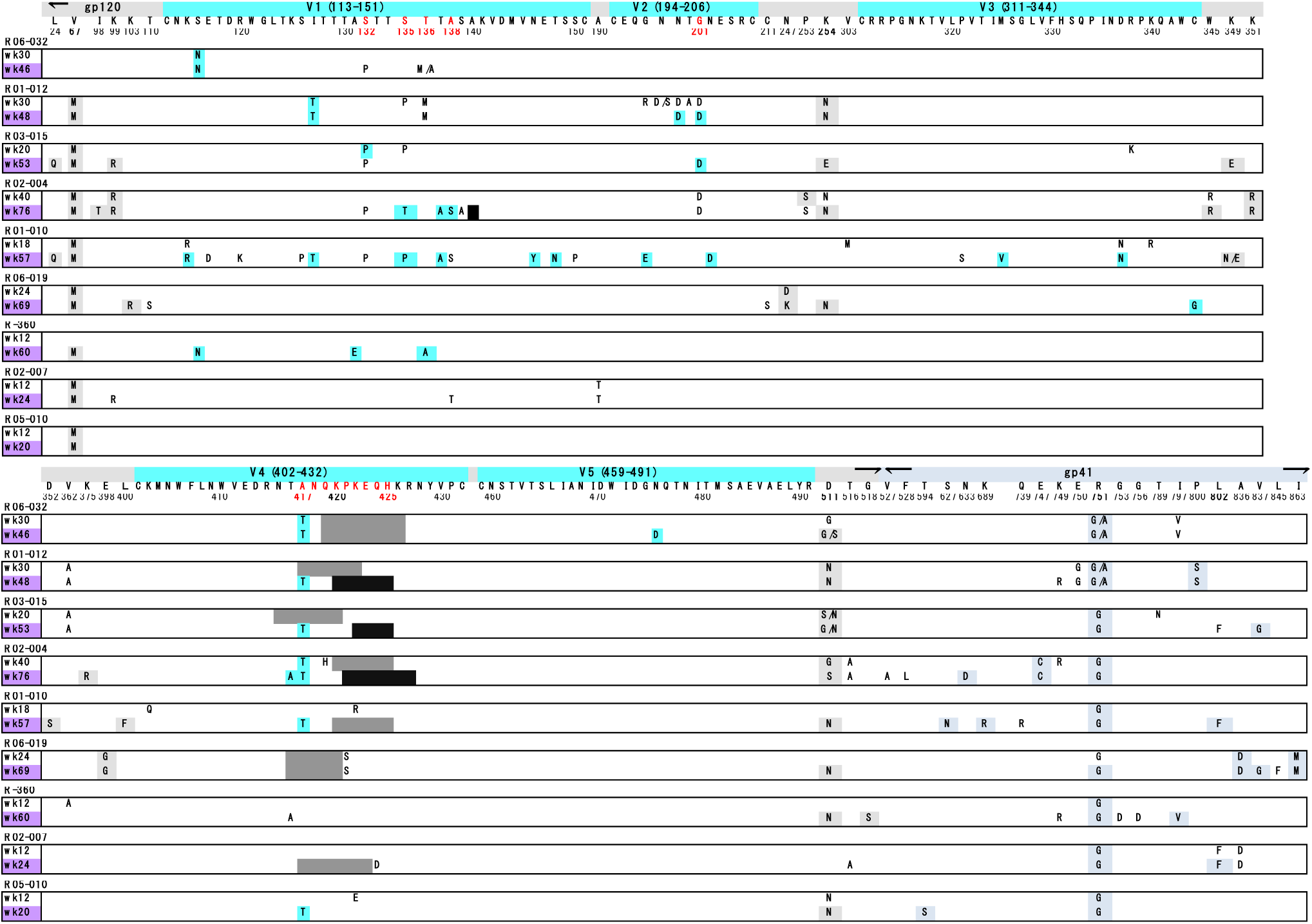
Env sequence variation pattern is not a major characteristic of NAb induction. SIV_mac239_ Env sequence variations before and after NAb induction. For each animal, the upper time point shows pre-induction and the lower time point (in purple) shows post-induction (first or second time point with NAb titer > 1:2). Dominant residue changes are shown as follows: gp120 (gray), variable regions V1-V5 (blue), and gp41 (blue gray). Subdominant mutations are uncolored. Unlabeled gray and black shadings show subdominant and dominant deletion of residues. Parental SIV_mac239_ sequence is listed for V1-V5. Residues mutated in multiple animals within V1-V5 are shown in red.

**Fig. 3-Supplement 1.**
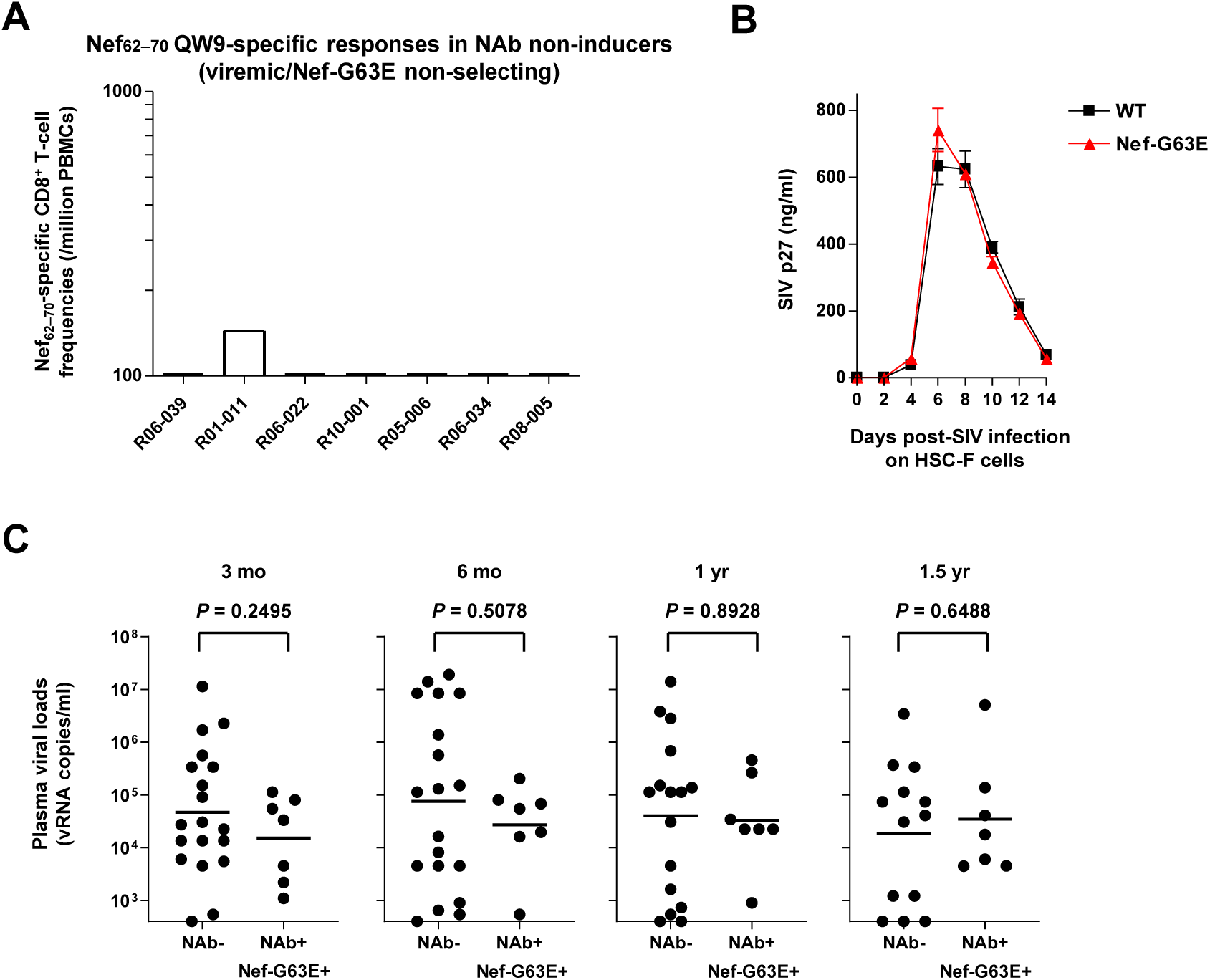
Selection of Nef-G63E mutant SIV is linked with Nef_62-70_-specific CD8^+^ T-cell response positivity. **(A)** Nef_62-70_-specific CD8+ T-cell responses around week 20 post-SIV challenge in seven viremic NAb non-inducers possessing either of the MHC-I alleles Mamu-B*039:01, Mamu-B*004:01 or Mamu-A1*032:02. **(B)** Temporal supernatant SIV p27 concentrations after wild type (WT) or Nef-G63E mutant SIV infection at multiplicity of infection (MOI) 0.001 on HSC-F cells. Data represent one of two independent experiments performed in triplicate. Bars: mean ± SEM. **(C)** Comparison of plasma viral loads in Nef-G63E-selecting NAb non-inducers (*n* = 7) versus non-inducers at indicated time points post-SIV challenge. Bars represent geometric means. log10-transformed values are compared by unpaired t tests.

**Fig. 4-Supplement 1.**
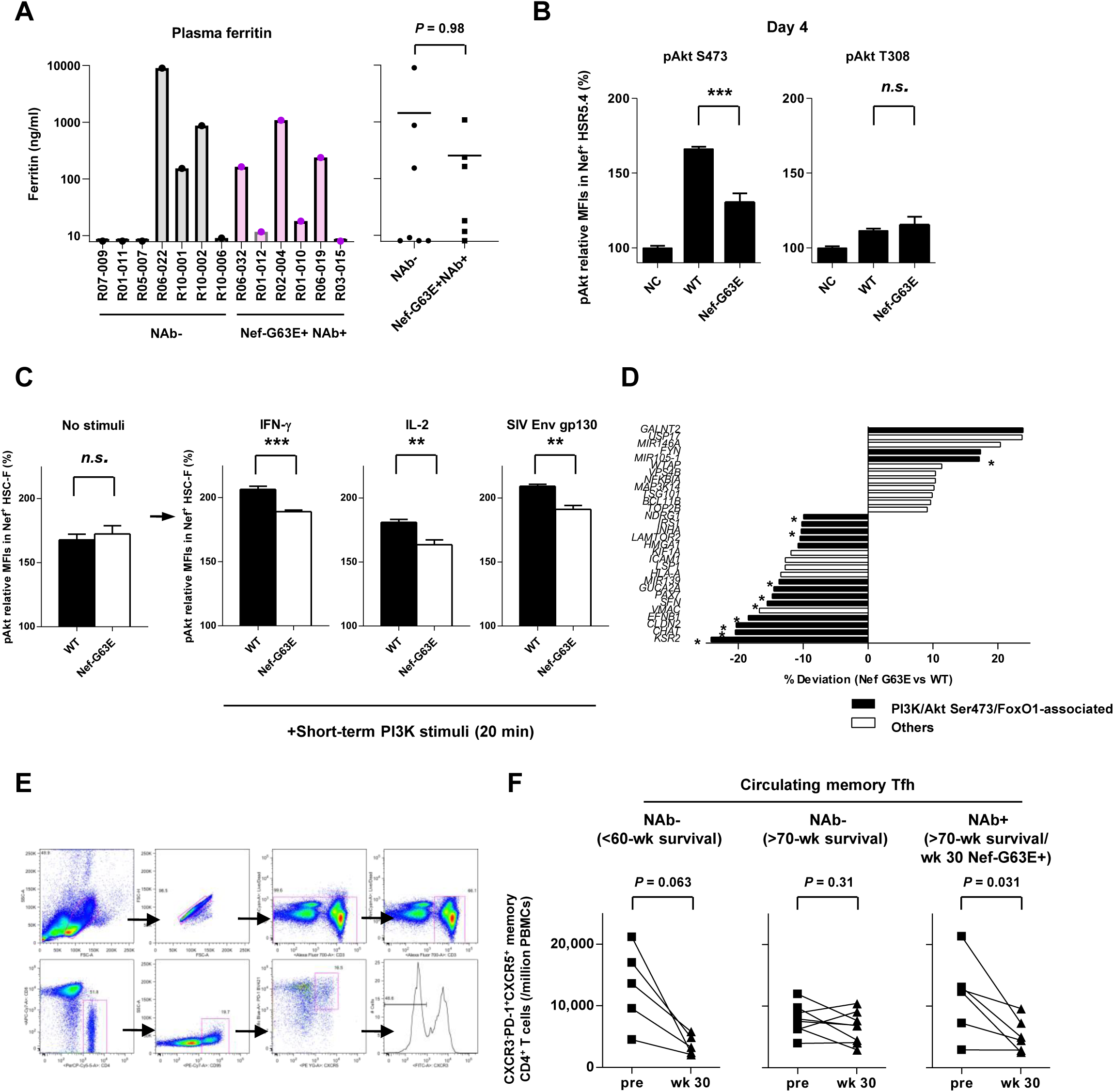
Decreased Akt hyperactivation properties of Nef-G63E mutant SIV. **(A)** Plasma ferritin levels (ng/ml) in examined NAb non-inducers versus Nef-G63E mutant SIV-selecting NAb inducers around year 1 post-infection. Bars represent mean in each group. Log-transformed values compared by unpaired t test. **(B)** Relative levels of pAkt Ser473 (left panel) and threonine (Thr) 308 (right panel) in SIV Nef^+^ HSR5.4 cells (MOI 0.1, 4 days p.i.). **(C)** Relative pAkt Ser473 levels in Nef^+^ HSC-F cells without stimuli and transiently pulsed with IFN-γ, IL-2 or SIV_mac_ Env gp130 after infection (MOI 0.2, 1 day). **(D)** Gene expression significantly differing between WT and Nef-G63E SIV infection (MOI 5, 1 day). Black bars show potentially Akt-related genes. Asterisks show dynamics concordant with the Nef-G63E phenotype (PI3K-pAkt Ser473 downstream downregulation or inhibition upregulation). **(E)** Representative flow cytometric gating (NAb inducer R01-012, wk 30 p.i.) of peripheral blood CXCR3^-^CXCR5^+^PD-1^+^ memory follicular CD4+ T cells. Cells were gated as CD3^+^CD8^-^ CD4^+^CD95^+^CXCR5^+^PD-1^+^CXCR3^-^ live singlet PBMCs. Available samples of six NAb inducers showing peak Nef-G63E mutant selection around week 30 p.i. and thirteen viremic NAb non-inducers were tracked. **(F)** Comparison of peripheral CXCR3^-^CXCR5^+^PD-1^+^ memory Tfh cell frequencies between pre-SIV infection and around week 30 post-infection in examined NAb non-inducers developing disease before week 60 (left: *n* = 5) and alive more than 70 weeks (middle: *n* = 8) and Nef-G63E-detected NAb inducers alive more than 70 weeks (right: *n* = 6). Analyzed by Wilcoxon signed-rank tests. MFIs relative to those in uninfected controls (%) are shown. **: *P* < 0.01, ***: *P* < 0.001, n.s.: not significant between WT and Nef-G63E mutant by unpaired t tests (B-C). Data represent one of two (B, C) independent experiments performed in triplicate (C: unstimulated) or quadruplicate (B, C: PI3K-pulsed). Bars: mean ± SEM.

**Fig. 4-Supplement 2.**
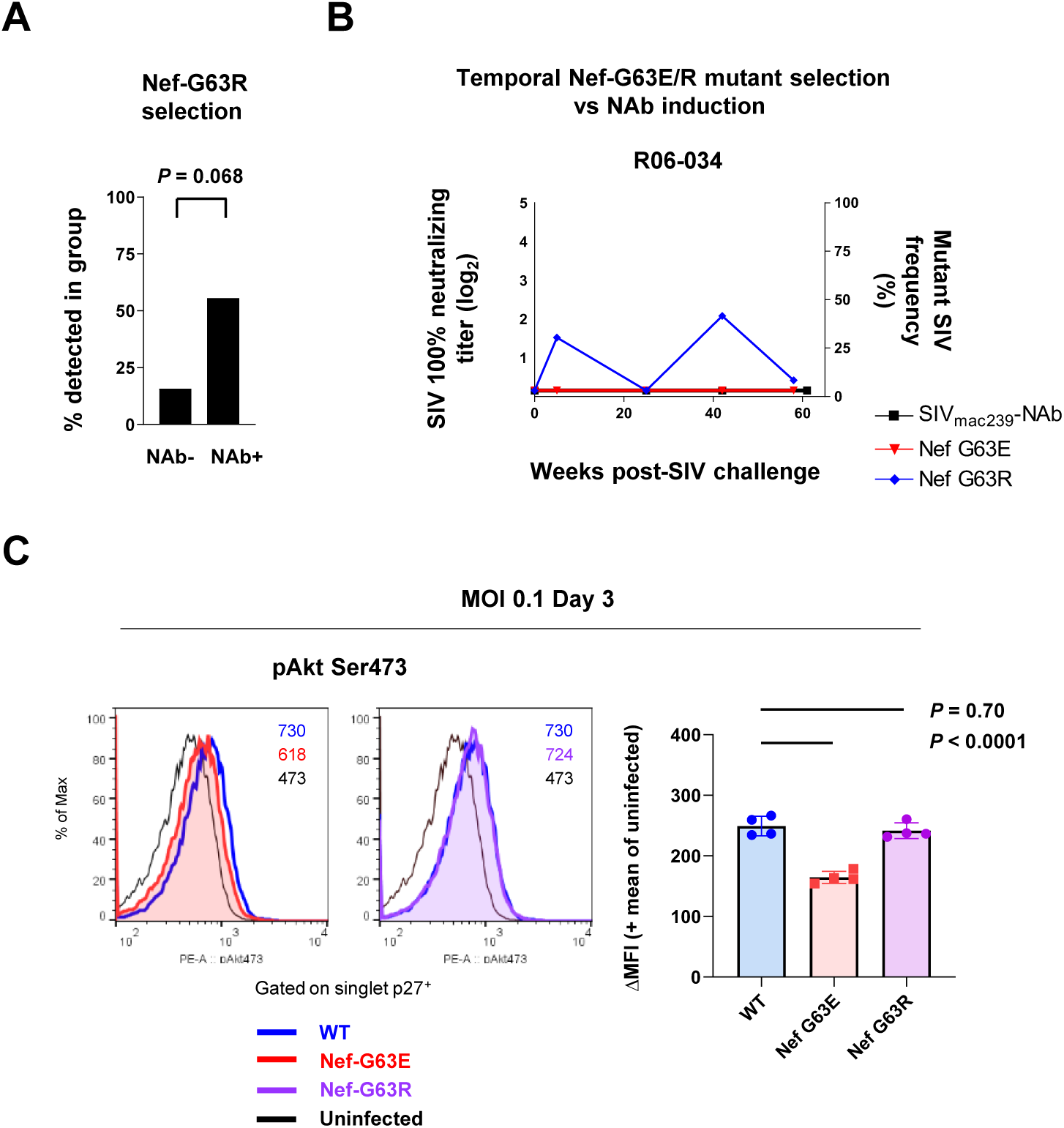
Nef-G63R mutant SIV properties. **(A)** Comparison of frequencies of macaques selecting for Nef-G63R mutant in plasma between NAb non-inducers and inducers. Compared by Fisher’s exact test. **(B)** Temporal relationship of Nef G63R frequencies in plasma virus in a NAb non-inducer macaque R06-034 preferentially selecting for Nef-G63R but not Nef-G63E mutant. Black boxes (left Y axis) show log_2_ NAb titers; red triangles and blue diamonds (right Y axis) show percentage of G63E and G63R mutations detected by subcloning (15 clones/point on average), respectively. **(C)** Left: representative histograms of relative pAkt serine (Ser) 473 levels in p27^+^ subpopulations after WT, Nef-G63E or Nef-G63R mutant SIV infection at MOI 0.1 on HSC-F cells. Numbers show pAkt Ser473 mean fluorescence intensities (MFIs) for each. Right: Deviation of pAkt Ser473 MFIs in p27^+^ HSC-F cells compared with mean MFI of uninfected cells. Analyzed by one-way ANOVA with Tukey’s post-hoc multiple comparison tests. Data represent one of two independent experiments performed in quadruplicate with similar results.

**Fig. 5-Supplement 1.**
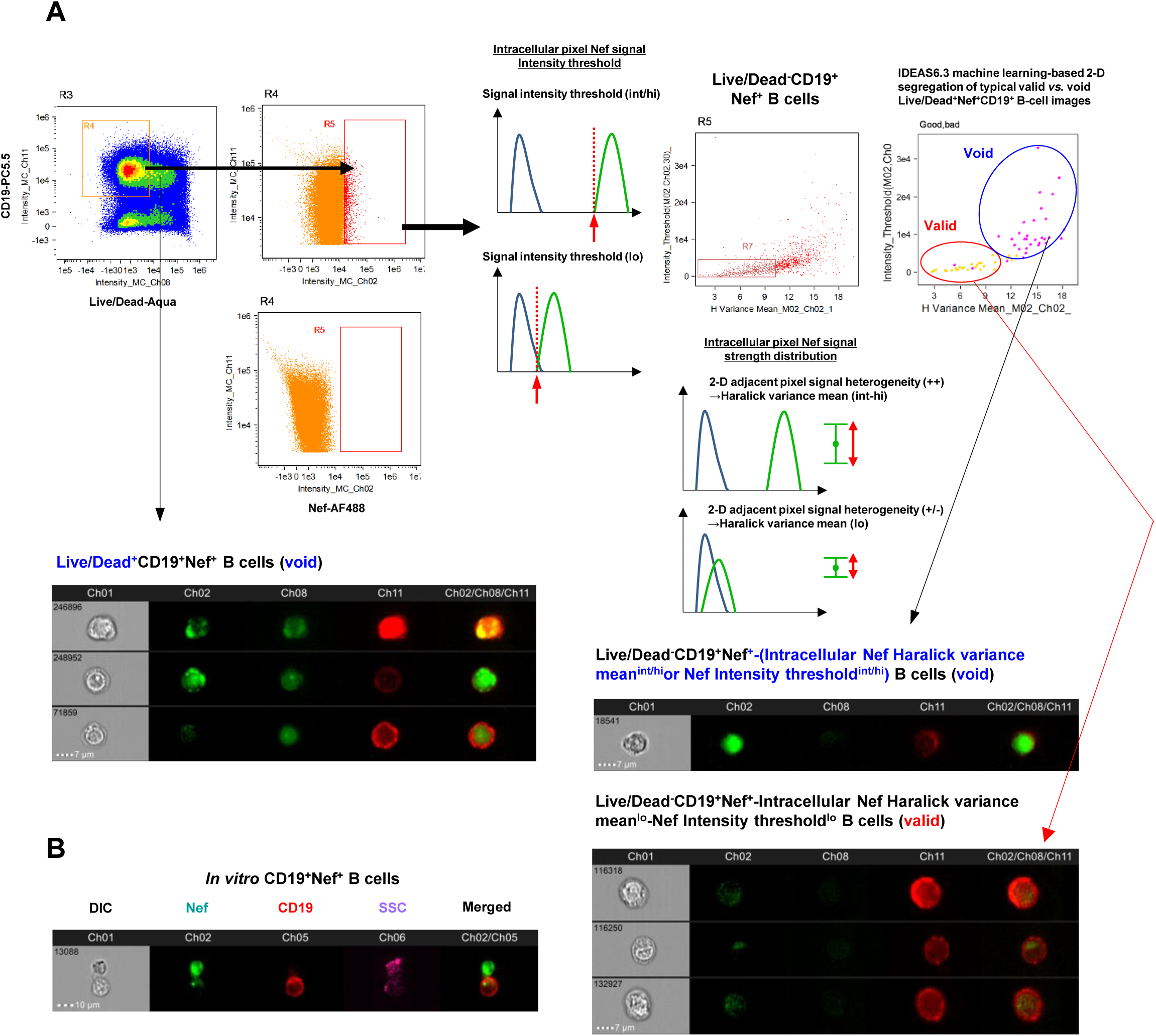
Machine learning-verified morphological gating of Nef-invaded B cells *in vivo*. **(A)** Representative gating strategy plus schematic of inguinal lymph node Nef^+^CD19^+^ B cells of an SIV_mac239_-infected rhesus macaque analyzed by ImageStreamX MK II imaging flow cytometry (R10-012, week 160 p.i.). First-step incorporation of Live/Dead staining gates out CD19^+^ B cells staining positive for Nef potentially due to pre-experimental membrane permeation (e.g., image #246896, #248952 and #71859). Second-step dual gating for intracellular Nef signal pixel Haralick variance mean (H Variance mean, X axis), a textural feature calculated from a gray level co-occurrence matrix (GLCM) for each image representing adjacent pixel signal strength heterogeneity and Nef signal intensity threshold (Y axis) gates out CD19^+^ B cells showing potentially non-specific, strong intracellular binary clustered staining pixels for Nef deriving a large variance (X axis), and/or cells stained overtly strong for Nef and hence becoming void for feasible image verification (high threshold) (Y axis), both likely originating stochastically from post-experimental rupturing procedures including membrane permeation (e.g. image #18541). This two-step noise cancellation results in acquisition of a Live/Dead^-^-Nef signal Haralick variance mean^lo^-Nef signal intensity threshold^lo^-Nef^+^CD19^+^ B-cell population (e.g. image #116318, #116250 and #132927) with a pericellular Nef^int-lo^ staining, biologically concordant with Nef membrane anchoring, resembling Nef^+^ cells experimentally generated *in vitro* and showed the highest score in IDEAS6.3 machine learning module scoring for segregation of typical void vs. valid images (upper right). This gating was implemented for counting and analysis of Env-specific B-cell Nef invasion *in vivo* (**Figure 5**). **(B)** Detection of contact-dependent Nef invasion from SIV-infected HSC-F CD4^+^ T cells to cocultured Ramos B cells *in vitro*. One typical image of an intercellular conduit (SSC, purple) with Nef (green) protrusions from infected HSC-F cells to CD19^+^ (red) Ramos B cells from two independent experiments are shown.

**Fig. 7-Supplement 1.**
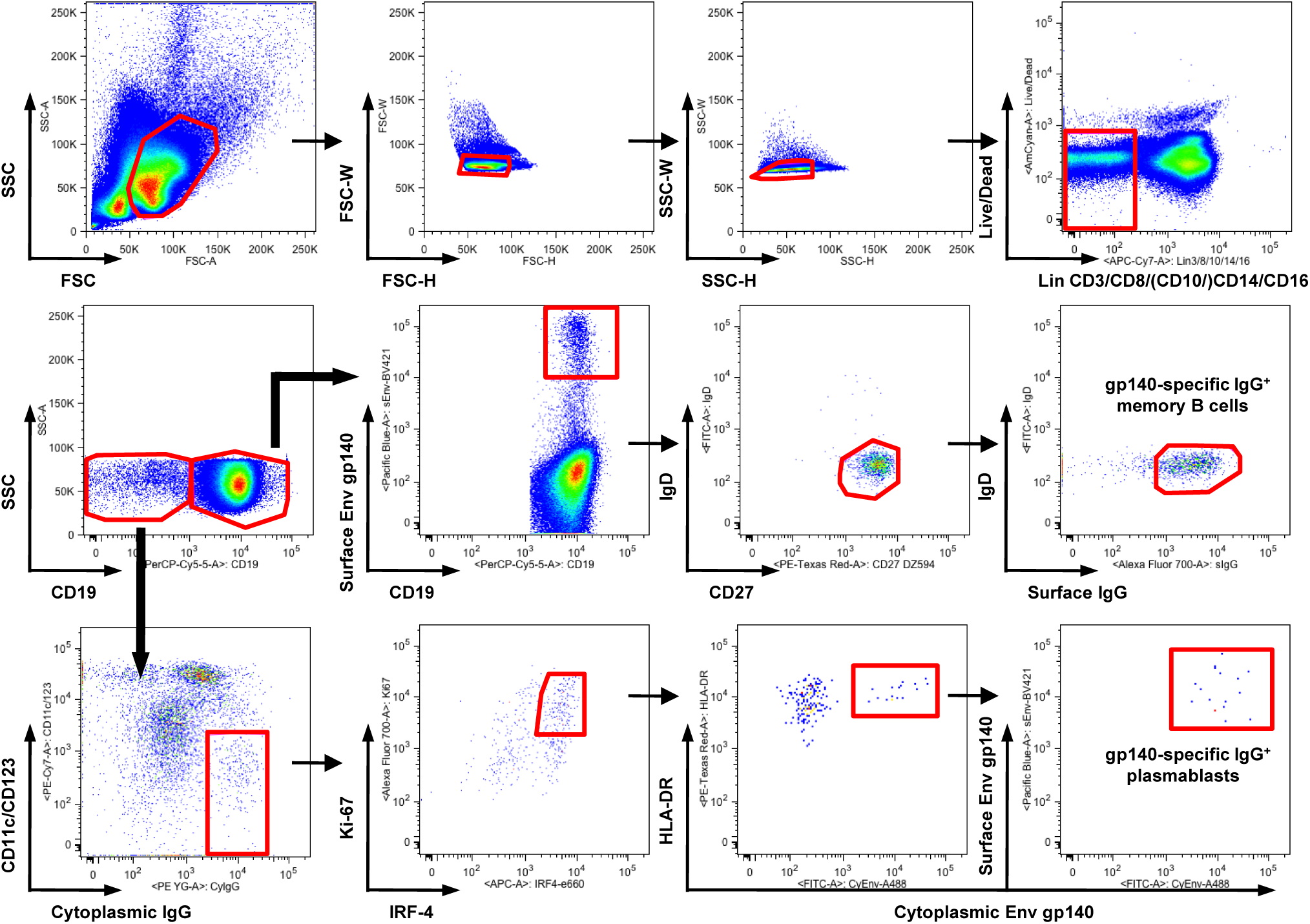
Full gating strategy of SIV_mac239_ Env gp140-specific B-cell responses. Full gating (R02-004, week 32 p.i.) of SIV Env gp140-specific memory B cells (B_mem_) and plasmablasts (PBs) shown in **Figure 7**.

## Key Resources Table

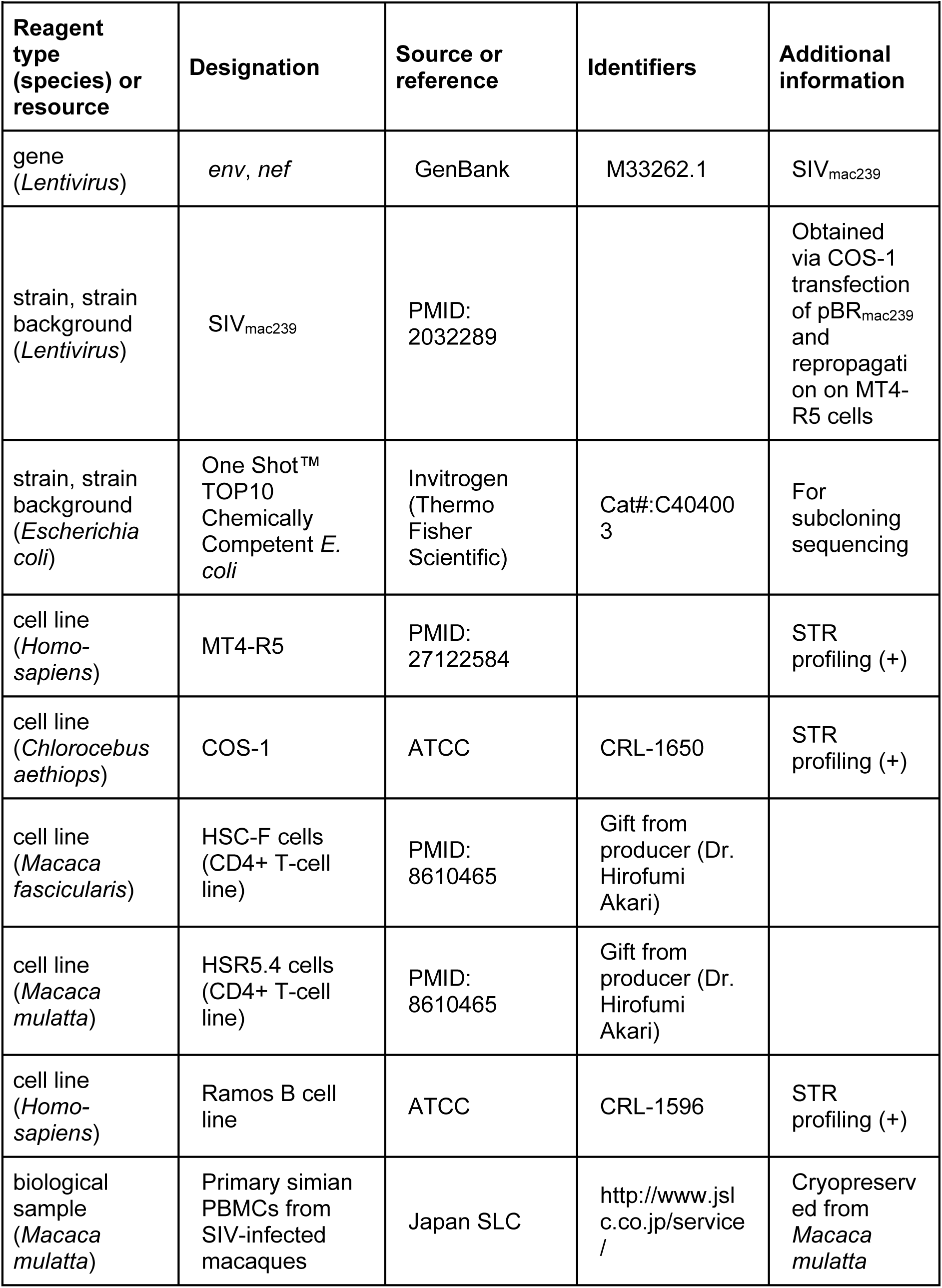
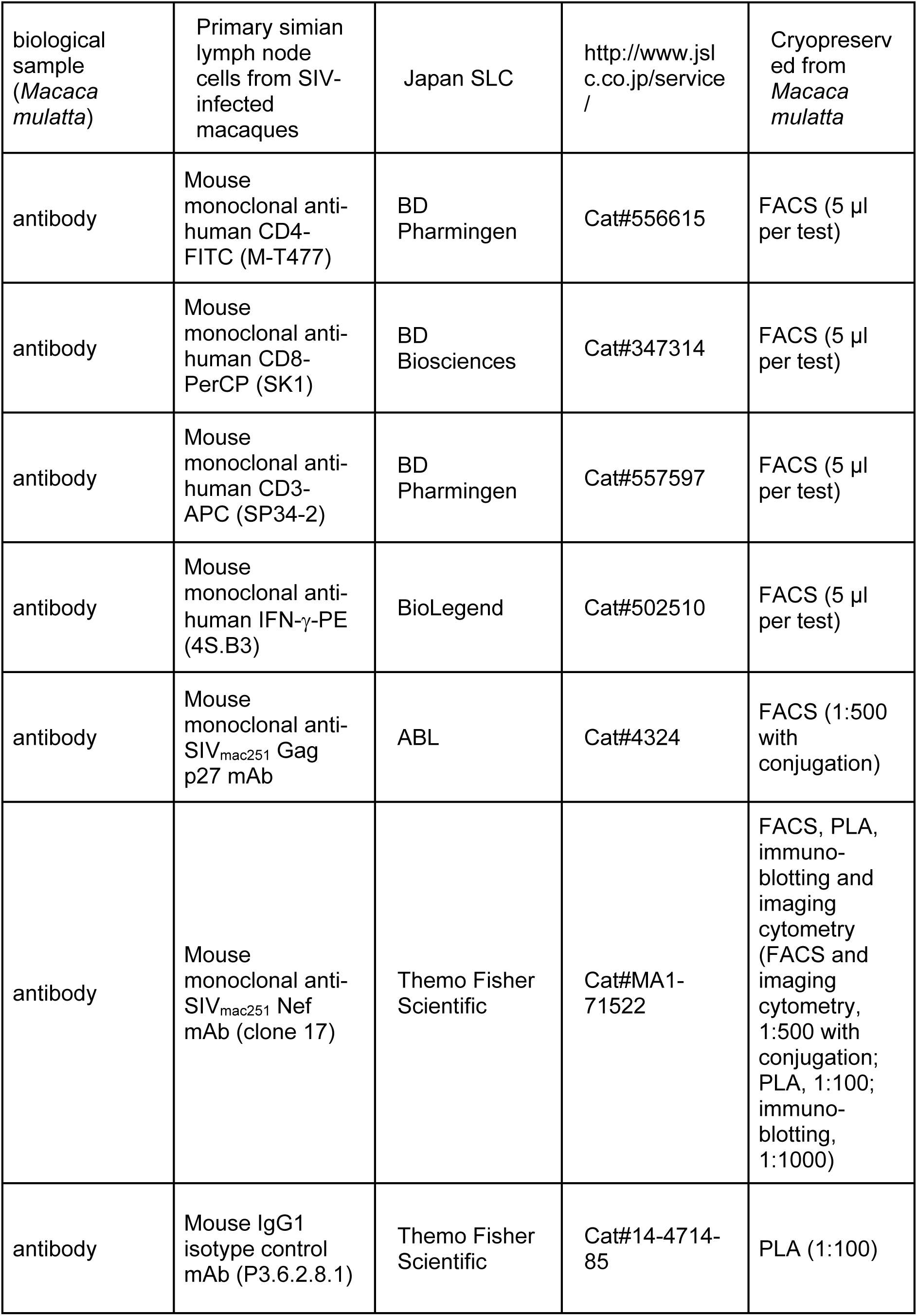
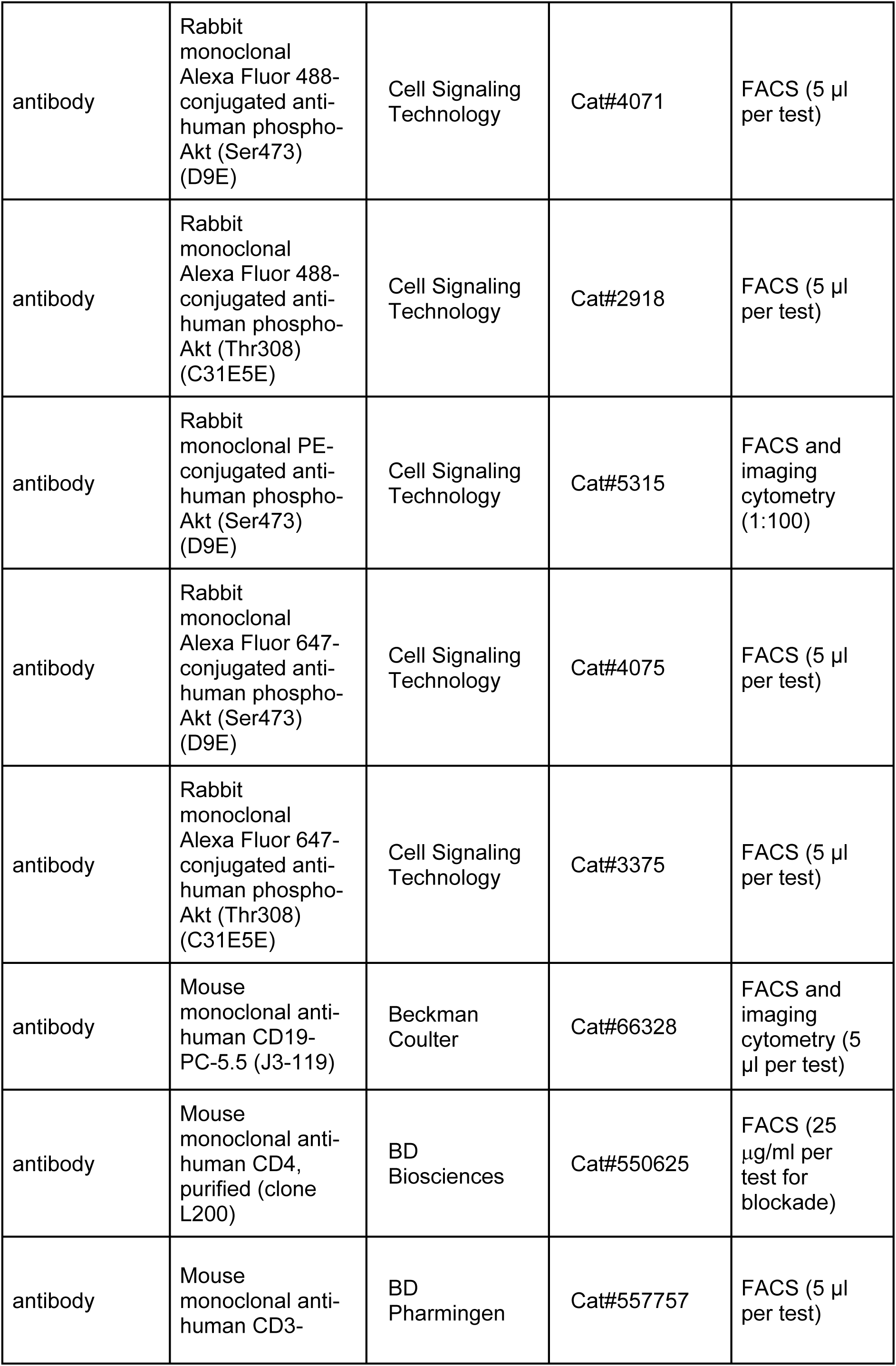
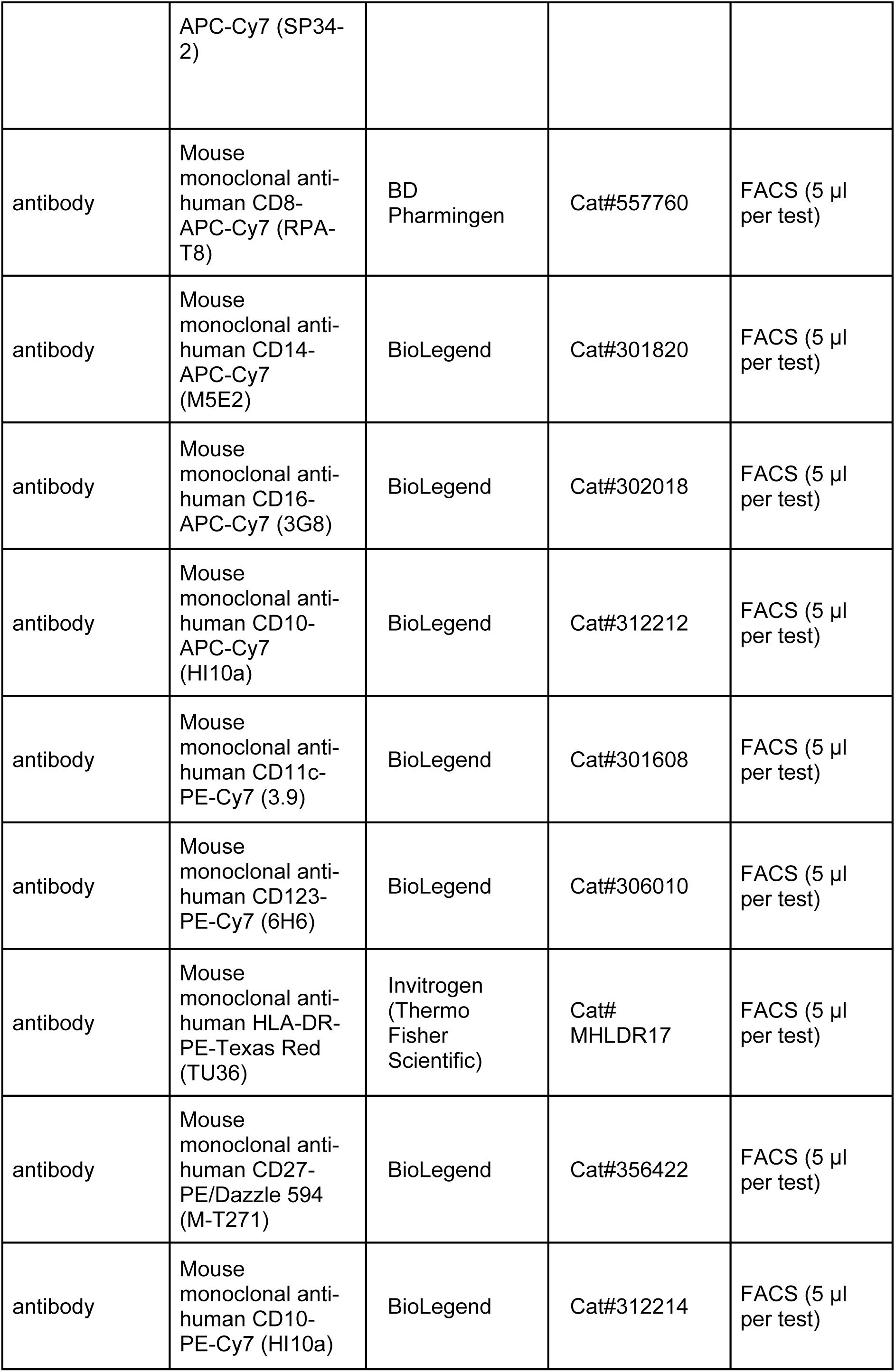
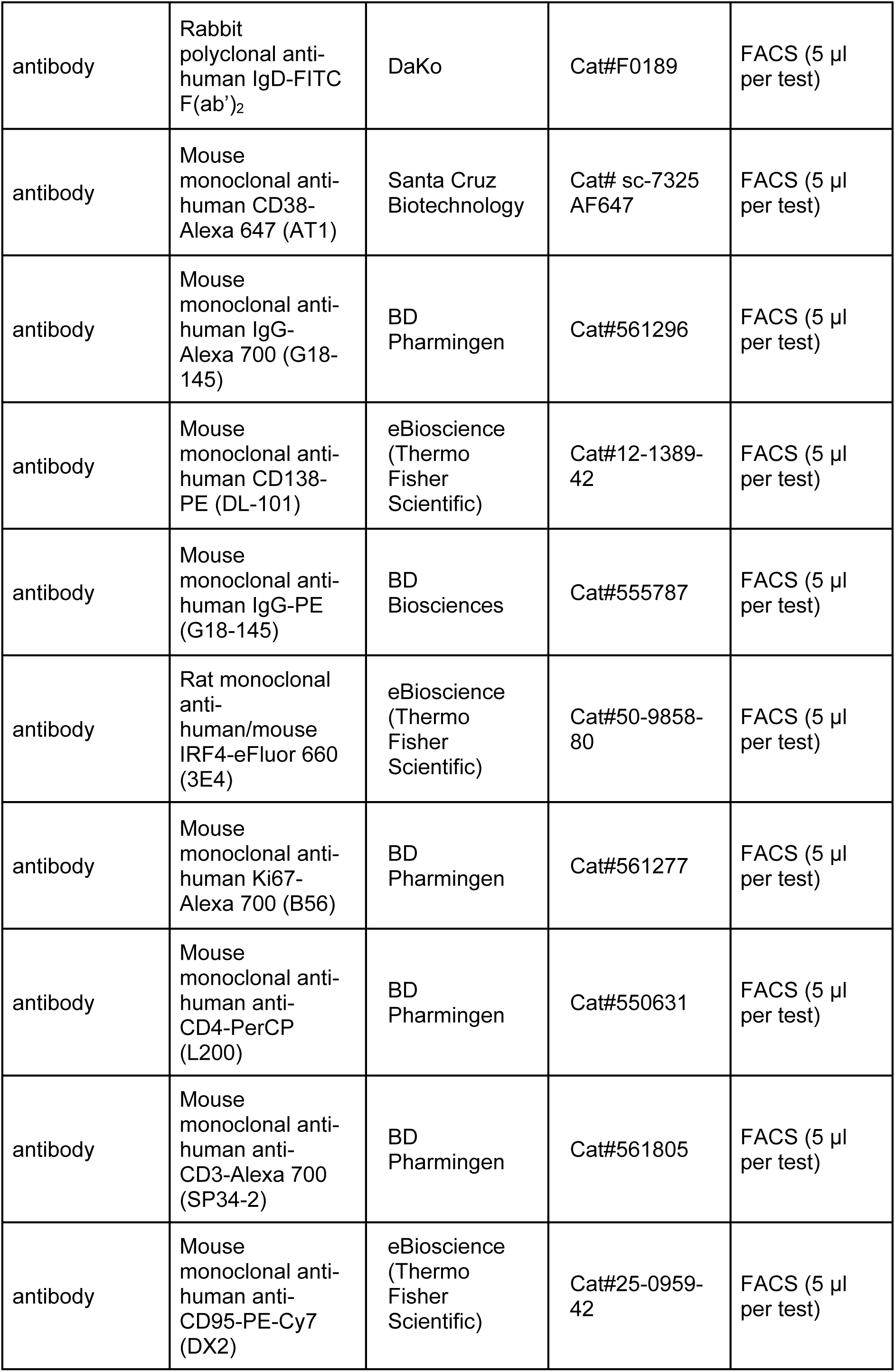
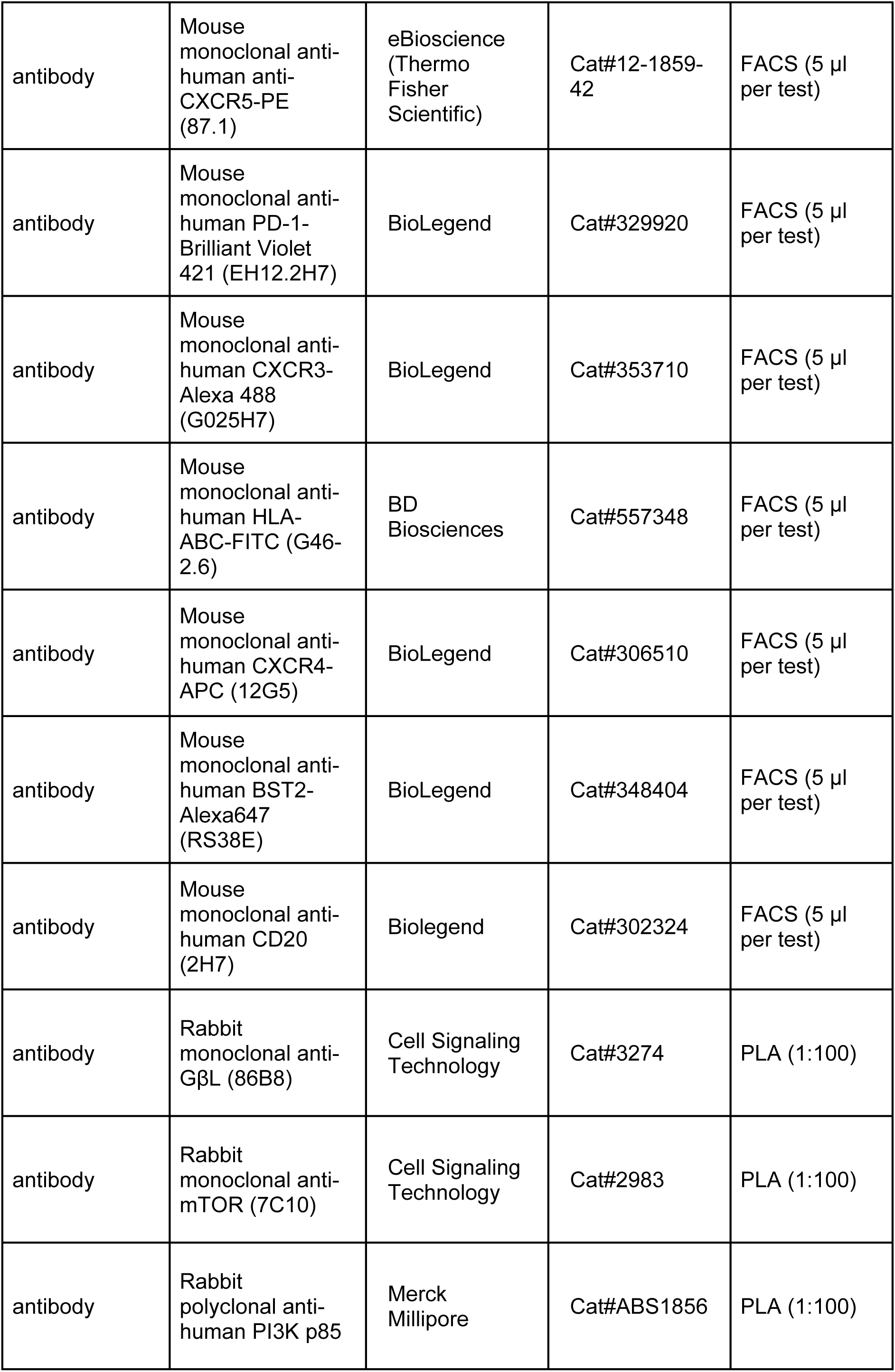
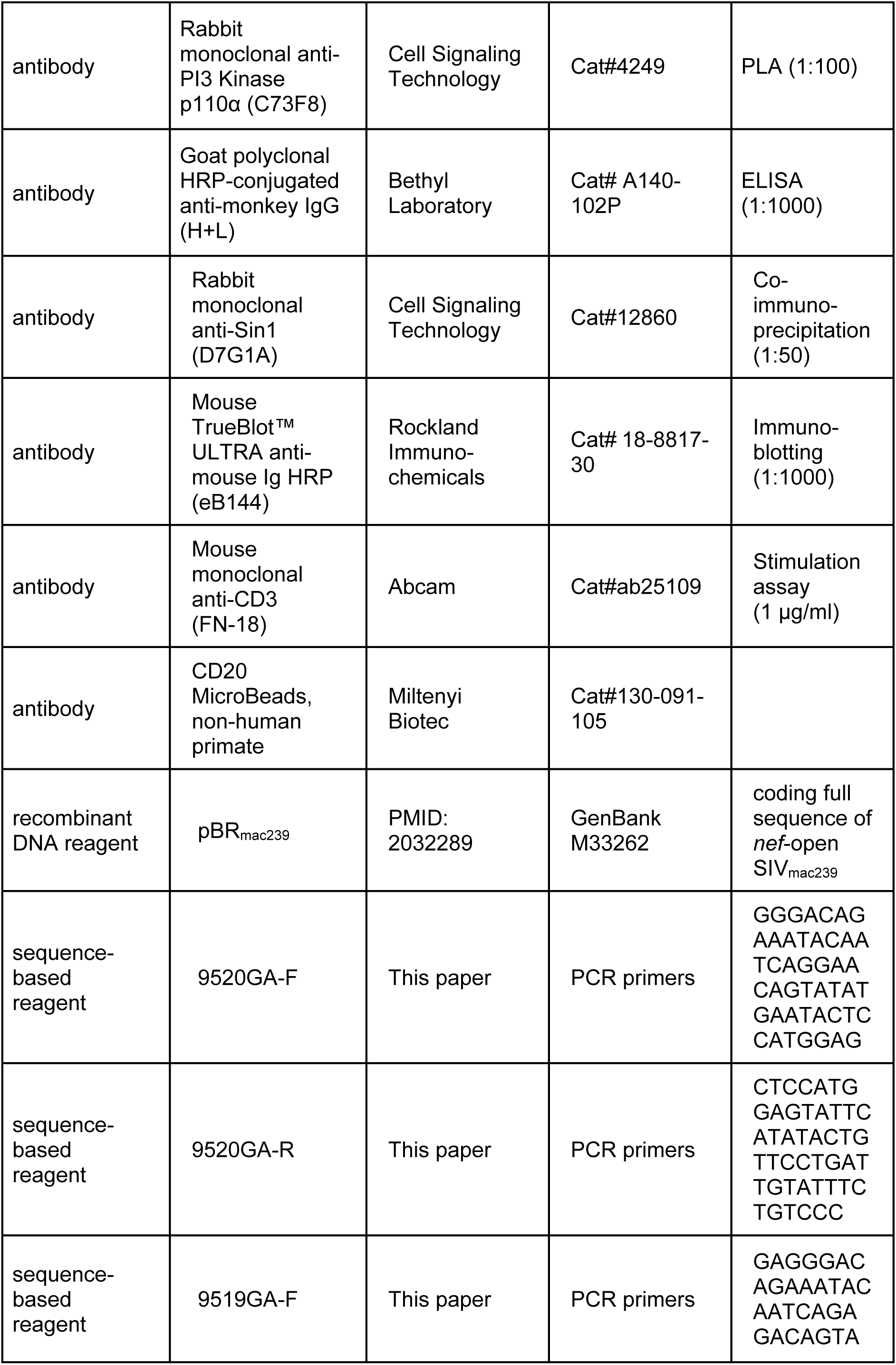
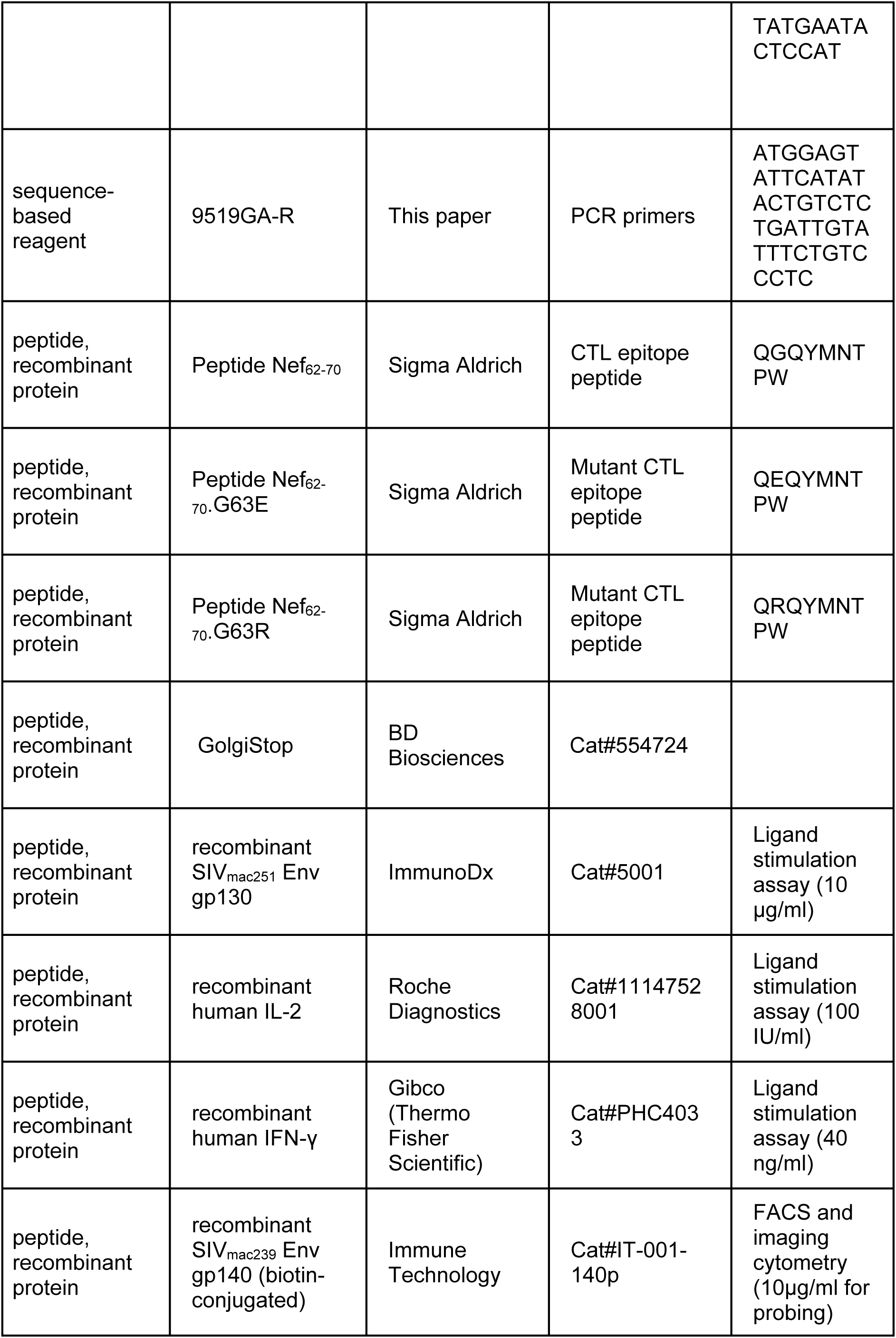
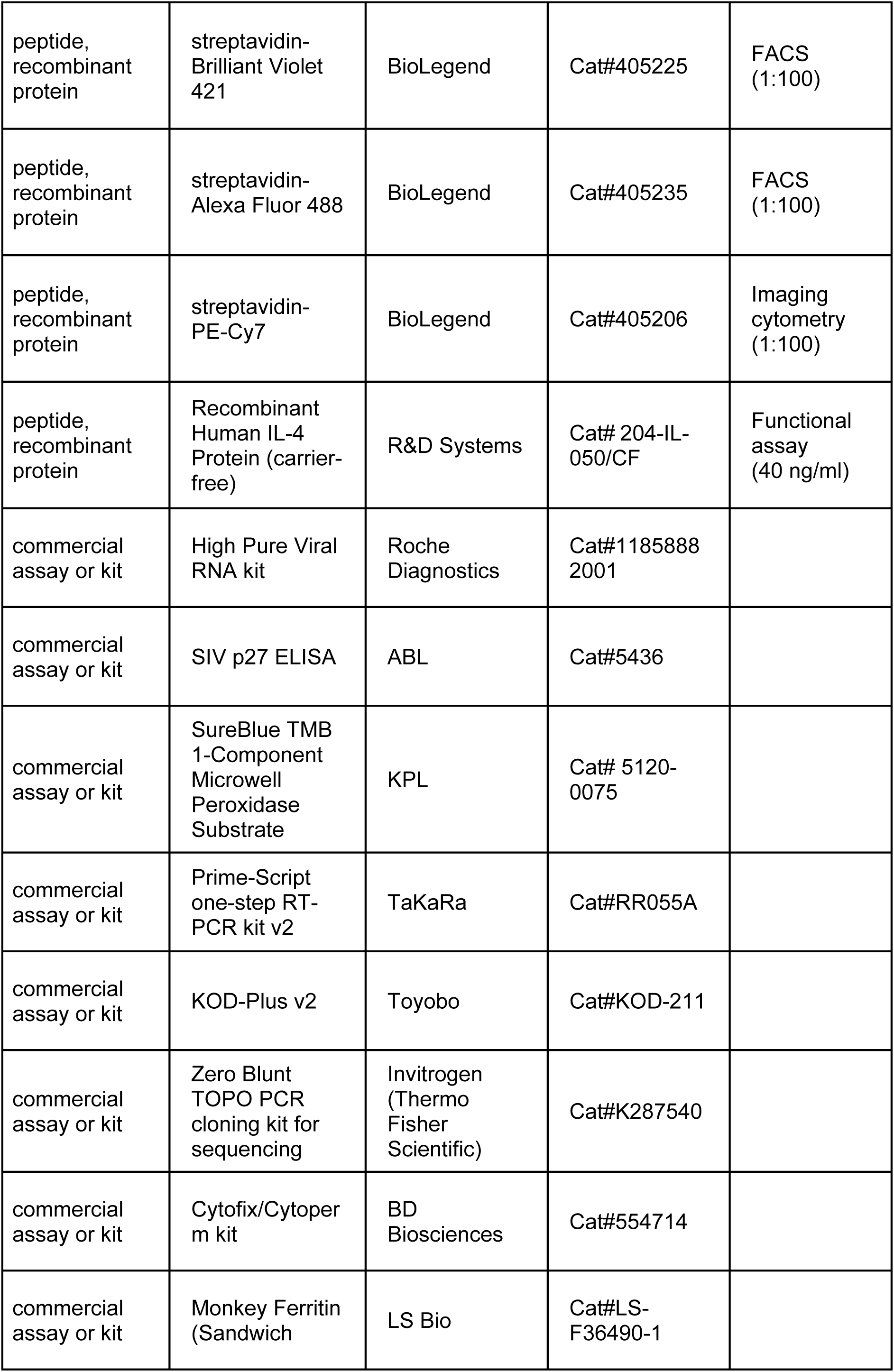
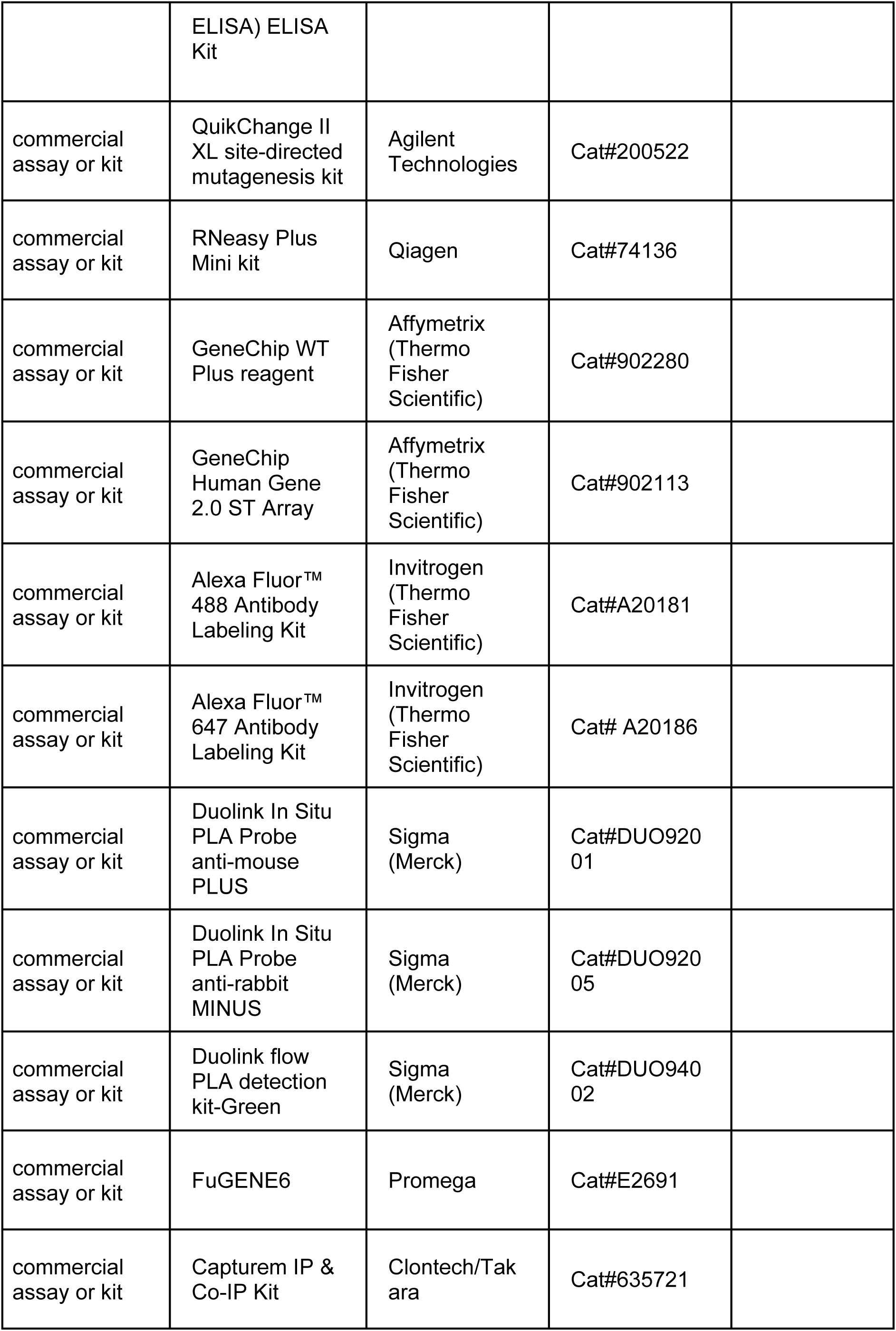
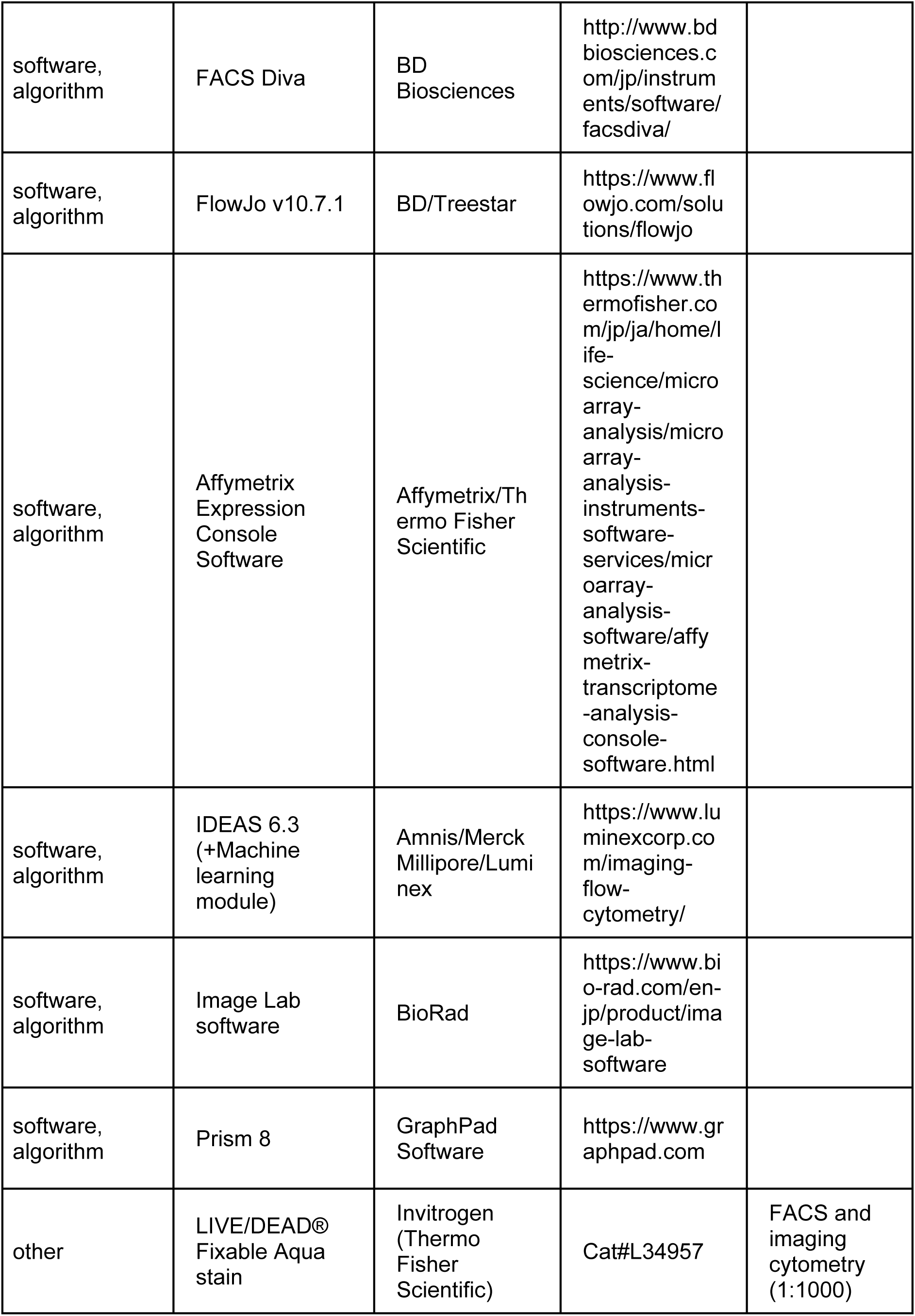
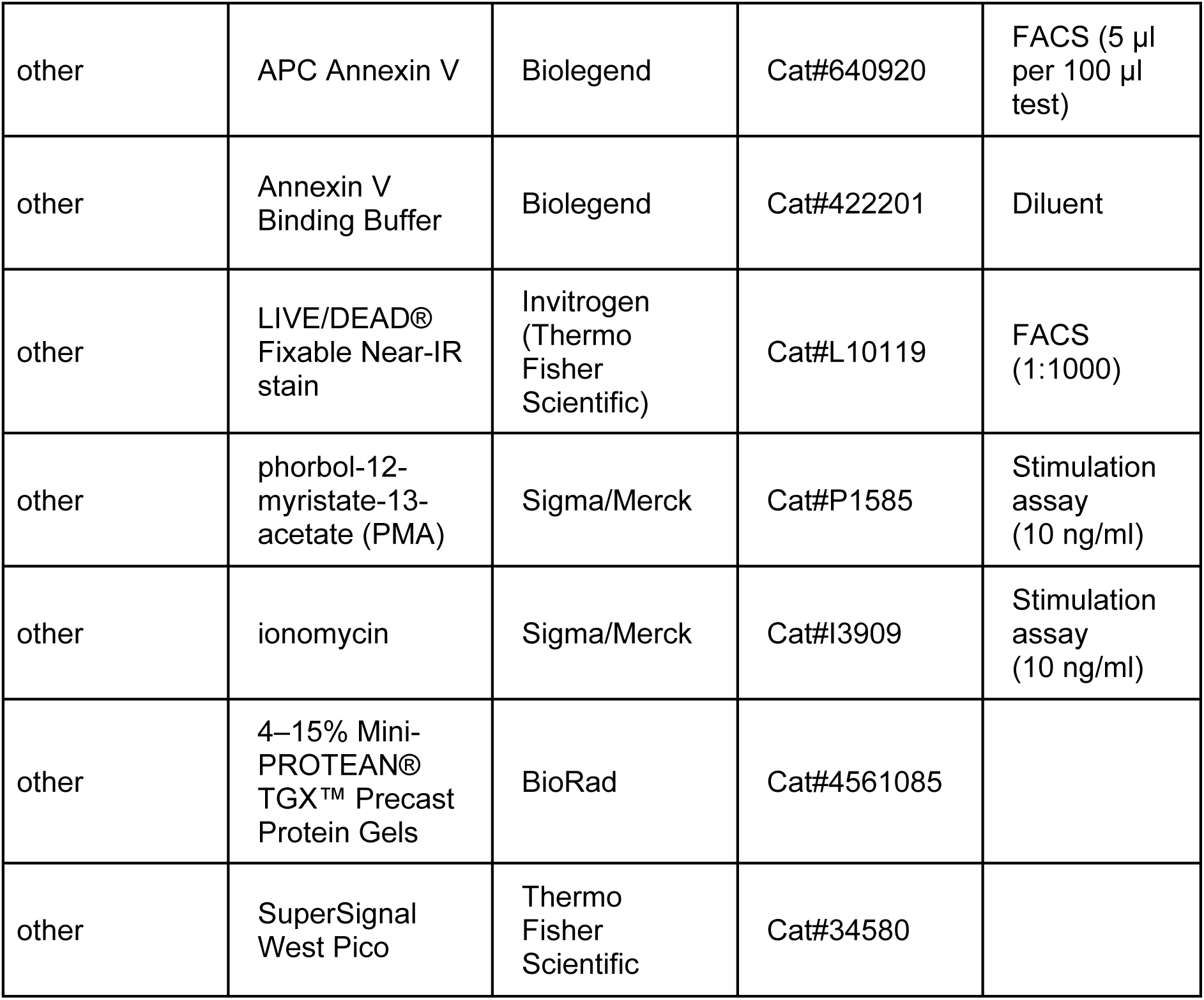

